# Multi-drug tolerance in *Leishmania* persister-like cells

**DOI:** 10.1101/2025.03.03.641139

**Authors:** Allison Aroni-Soto, Pieter Monsieurs, Odin Goovaerts, Kaoutar Choukri, Basudha Khanal, Wim Adriaensen, Jean-Claude Dujardin, Michael P. Barrett, Malgorzata A. Domagalska

## Abstract

Leishmaniasis is caused by parasitic protozoa of the genus *Leishmania* and is found widely across the tropics and sub-tropics, afflicting hundreds of thousands of people. The disease is notoriously difficult to treat. Here, we present evidence of the existence of persister-like cells in cultured *Leishmania* populations, induced upon exposure to normally lethal doses of antimony, a widely used anti-leishmanial. Persisters are a small fraction of non-proliferative cells with reduced metabolism that are adapted to withstand a variety of environmental assaults, including lethal doses of antimicrobials. We show that *Leishmania* persister-like cells survive lethal doses of antimonials by adopting a quiescence phenotype characterised by reduced proliferation, constrained metabolism, and diminished mitochondrial membrane potential. What is more, these cells demonstrate cross-tolerance to other anti-leishmanial drugs. Wild-type persister-like cells reverted to similar levels of drug susceptibility once the antimony-induced pressure was removed. Surprisingly, cells which had previously been selected for genetic changes causing resistance to antimony acquired a level of hyper-resistance after transient passage through the quiescent state, without further genetic change. Our results demonstrate the extreme versatility of this eukaryotic pathogen in adaptation to drug pressure and highlight the need for the development of new anti-leishmanials targeting non-proliferative forms.

## Main

The ability of microorganisms to resist the killing action of antimicrobial agents is emerging as one of the most serious public health threats and has been observed for antivirals, antibiotics, antifungals, and also antiparasitic compounds ^1^. Understanding the mechanisms that allow them to survive chemotherapy is essential. Several molecular mechanisms can underlie this phenomenon, but the most extensively studied one is drug resistance caused by acquired, heritable genetic changes, usually point mutations or insertion/deletion within genes in which these mutations enable cells to prevent the drug’s toxic capacity ^2,3^. Importantly, resistant pathogens remain metabolically active and can grow and replicate in the presence of drugs. However, microorganisms can also adopt a physiological state of transient quiescence, manifested by non-proliferation and reduced metabolism. This leads to non-heritable drug tolerance in cases where drug activity depends upon metabolic activities suppressed in quiescence ^4^. This can affect whole-microbial populations, but heterogeneity within microbial populations may allow the survival of a small number of specialized quiescent cells, termed ‘persisters’, in the presence of an antimicrobial agent ^5^. After the removal of drug pressure, persisters resume growth and switch back to their drug-sensitive phenotype. The presence of persisters is of particular importance as their quiescent phenotype associated with exposure to a single drug often leads to multidrug tolerance ^6–8^.

Quiescence is a non-proliferative, physiological resting state that can be adopted by many infectious microorganisms, including bacteria, fungi, and some protozoan parasites as a strategy to survive nutrient limitation, host immunity, and chemotherapy ^9^. This reversible state is characterized by an active and regulated cellular and molecular remodelling programme resulting in reduced metabolism, transcription, and translation ^9,10^. Importantly, while overall metabolism is switched into an energy-saving mode, crucial processes allowing the survival of the cells under stress conditions remain active, and some processes are even selectively upregulated in the quiescent state ^11^.

*Leishmania* are digenetic protozoan parasites that cycle between the sand fly vector and mammalian host. They cause a spectrum of diseases, the most severe form being visceral leishmaniasis (VL), which is lethal if left untreated. *Leishmania donovani* is the etiological agent of VL in the Indian subcontinent (ISC) and in East Africa. Antimonials (Sb) were the first drugs used in the treatment of VL, but they have been replaced first by miltefosine and later by Amphotericin B (AmB) in the ISC due to loss of efficacy attributed to the emergence of drug resistance ^12–15^. Phylogenomic analysis of 204 *L. donovani* isolates, followed by experimental studies allowed the identification and characterization of different genomic pre-adaptations to antimonials in *Leishmania* in the ISC ^16^. They identified i) a small group of ISC1 parasites, which diverged early and have no obvious genomic pre-adaptation to antimony, ii) a dominant, so-called core group of parasites consisting of several closely related lineages which are pre-adapted to antimony through either the presence of the intra-chromosomal amplification of the H-locus, which contains the MRPA gene, a known driver of antimony resistance or iii) another group with increased H-locus copy number, plus a 2-nt insertion in the AQP1 gene (ISC5), that encodes a key importer of antimony whose loss also contributes to resistance to the drug ^16,17^.

In terms of drug responses in *Leishmania,* most research has focused on identification of potential drug resistance signatures in natural populations and in experimental evolution studies ^16,18–21^. However, recent evidence points to the existence of quiescence and its role in adaptation to drug pressure ^22–26^. Quiescence in *Leishmania* was inferred in mice when amastigotes of *L. mexicana* were shown to have a very low replication rate, together with low rates of RNA synthesis, protein turnover, and membrane lipid synthesis ^27^. Another mouse study revealed heterogeneity among *L. major* amastigotes with non-replicating parasites distinguished from slowly replicating populations ^28^. Another study of *L. mexicana* in mice revealed quiescent populations within collagen-rich mesothelium surrounding lesions, where miltefosine accumulation was also limited ^25^. Using an in vitro antimony-exposure model, we showed that *L. lainsoni* parasites exhibit hallmarks of a quiescence state and that drug-tolerant and persister-like cells of genetically diverse strains of *L. braziliensis* could be detected under high doses of potassium antimonium tartrate (PAT) ^23,24^. Recently, Dirkx and colleagues showed that long-term hematopoietic stem cells provide a parasite niche during treatment failure in an animal model of VL infection ^29^. Bone marrow stem cell-associated parasites were proposed to enter into quiescence after 2-3 replication cycles with parasites emerging post-quiescence with increased apparent infectivity ^26^.

In selecting drug resistance in laboratory studies in *Leishmania*, it has become clear that these organisms have particularly plastic genomes and hundreds of karyotypes can emerge from a single clonal population grown in culture ^17,21,30–32^. This phenomenon, termed mosaic aneuploidy, is postulated to underlie phenotypic heterogeneity and facilitate adaptation to changing environmental conditions, including drug pressure ^32,33^. Generally, persisters in bacteria do not emerge due to genetic changes, however, it is important to ascertain whether a similar process relates to quiescence in *Leishmania* in the selection of drug-tolerant persister-like cells.

In this study, we addressed: i) the role of quiescence and population heterogeneity in the survival of *L. donovani* promastigotes under normally lethal doses of PAT, and ii) whether the presence of genomically defined pre-adaptations to antimony has an impact on the quiescence and persister phenotypes under PAT exposure. We found compelling evidence that quiescence is a universal survival mechanism adapted by genomically distinct *L. donovani* strains. However, the impact of the transition through quiescence, on drug susceptibility differed, depending on the presence of genomic pre-adaptations to antimony. In non-preadapted parasites, the antimony tolerance phenotype was reversible in non-drug conditions. In contrast, in resistant parasites a small sub-population of persisters gave rise to a population of parasites with further increased resistance to antimony. The quiescent state observed in both types of persister-like cells can provide tolerance to PAT, but also other antileishmanial compounds. With this study, we demonstrate for the first time multidrug tolerance in persister-like cells of a eukaryotic pathogen.

## Results

### Isogenic *L. donovani* lines exposed to normally lethal doses of PAT exhibit intrinsic population heterogeneity in their drug susceptibility

To address whether heterogeneity within populations and quiescence plays a role in response to drug pressure in *L. donovani*, we developed an experimental model, in which we focused on the detection of a small subpopulation of surviving persister-like cells. Specifically, we aimed to identify cells that i) constituted a small subpopulation within isogenic lines, ii) could survive in the presence of a normally lethal dose of a drug and iii) showed a reduction in growth and metabolism associated with quiescent state when compared with the majority of cells. We chose trivalent antimony (PAT), as we showed previously that quiescent cells could be observed in the presence of high doses of this drug in a dermotropic species of *Leishmania*, *L. braziliensis* ^23^.

We selected ten cloned strains of *L. donovani* isolated in the ISC, with different levels of genomic pre-adaptation to antimony: i) three strains where no obvious genomic changes indicative of pre-adaptation to PAT resistance are known to be present (ISC1); these strains are highly susceptible to PAT; ii) seven strains belonging to different genetic groups (ISC4, 6, and 9) characterized by possessing one known genomic pre-adaptation to PAT, the intra-chromosomal amplification of the H-locus, which contains the MRPA gene, a known driver of antimony resistance; these strains show decreased susceptibility to PAT; iii) one strain with two known genomic pre-adaptations to PAT: a 2-nt insertion in the AQP1 locus, proposed to be associated with antimonial resistance observed in the ISC5 genotype in addition to the intra-chromosomal amplification of the H-locus; the strain shows a low susceptibility to PAT ^16,17^. We also included a reference *L. donovani* strain isolated in East Africa (LdBob), highly susceptible to antimony (Table 1) ^34,35^. Of note, one of the strains belonging to ISC6 group, BPK085 also carries a different 2-nt insertion mutation in the AQP1 locus, which results in a frameshift ^16^.

**Table 1.**
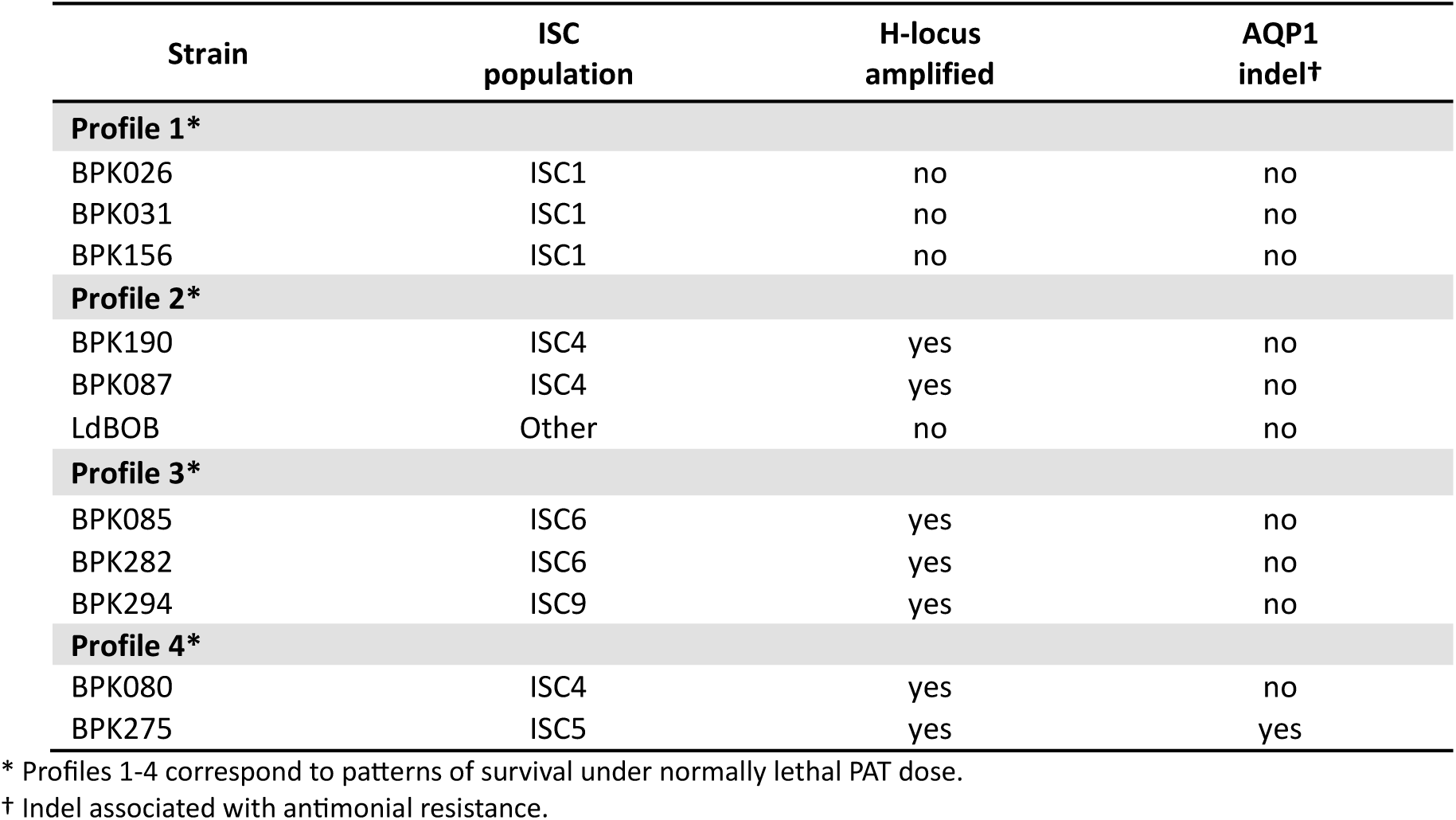
Cloned strains of *L. donovani* isolates from the Indian Subcontinent and reference strain (LdBOB) used in this study.

| Strain | ISC population | H-locus amplified | AQP1 indel† |
| --- | --- | --- | --- |
| **Profile 1\*** |  |  |  |
| BPK026 | ISC1 | no | no |
| BPK031 | ISC1 | no | no |
| BPK156 | ISC1 | no | no |
| **Profile 2\*** |  |  |  |
| BPK190 | ISC4 | yes | no |
| BPK087 | ISC4 | yes | no |
| LdBOB | Other | no | no |
| **Profile 3\*** |  |  |  |
| BPK085 | ISC6 | yes | no |
| BPK282 | ISC6 | yes | no |
| BPK294 | ISC9 | yes | no |
| **Profile 4\*** |  |  |  |
| BPK080 | ISC4 | yes | no |
| BPK275 | ISC5 | yes | yes |

\* Profiles 1-4 correspond to patterns of survival under normally lethal PAT dose.
† Indel associated with antimonial resistance.

By using this collection of natural parasite isolates with different levels of genetic pre-adaption to antimony we aimed to study the potential interplay between different heritable, genetically defined forms of drug resistance and quiescence in parasite survival under PAT pressure. We generated rEGFP lines for each of these strains to monitor transcriptional activity of rDNA locus as a negative biomarker of quiescence during exposure to PAT ^36^.

In the first step, we exposed promastigotes of all lines to a range of PAT concentrations to determine for each strain, the lowest concentration (Supp. Fig.1; Supp. Table 1) at which no metabolic activity could be detected in a resazurin based assay measured at the level of the whole population ^37^. By identifying normally lethal doses of PAT specific for each strain, we aimed to establish comparable experimental conditions for each strain in which physiologically quiescent-like cells would be observed, as shown for bacteria. We then subjected each line to the respective strain-specific normally lethal PAT doses over 10 days (with PAT application repeated at day 5), and we monitored viability and growth (Supp. Table 2; Supp. Fig. 2) and established the time-kill curves (Fig.1; Supp. Fig.3) during the exposure. Overall, we identified four broad viability profiles in response to PAT pressure (Fig.1; Supp. Table 2).

**Figure 1.**
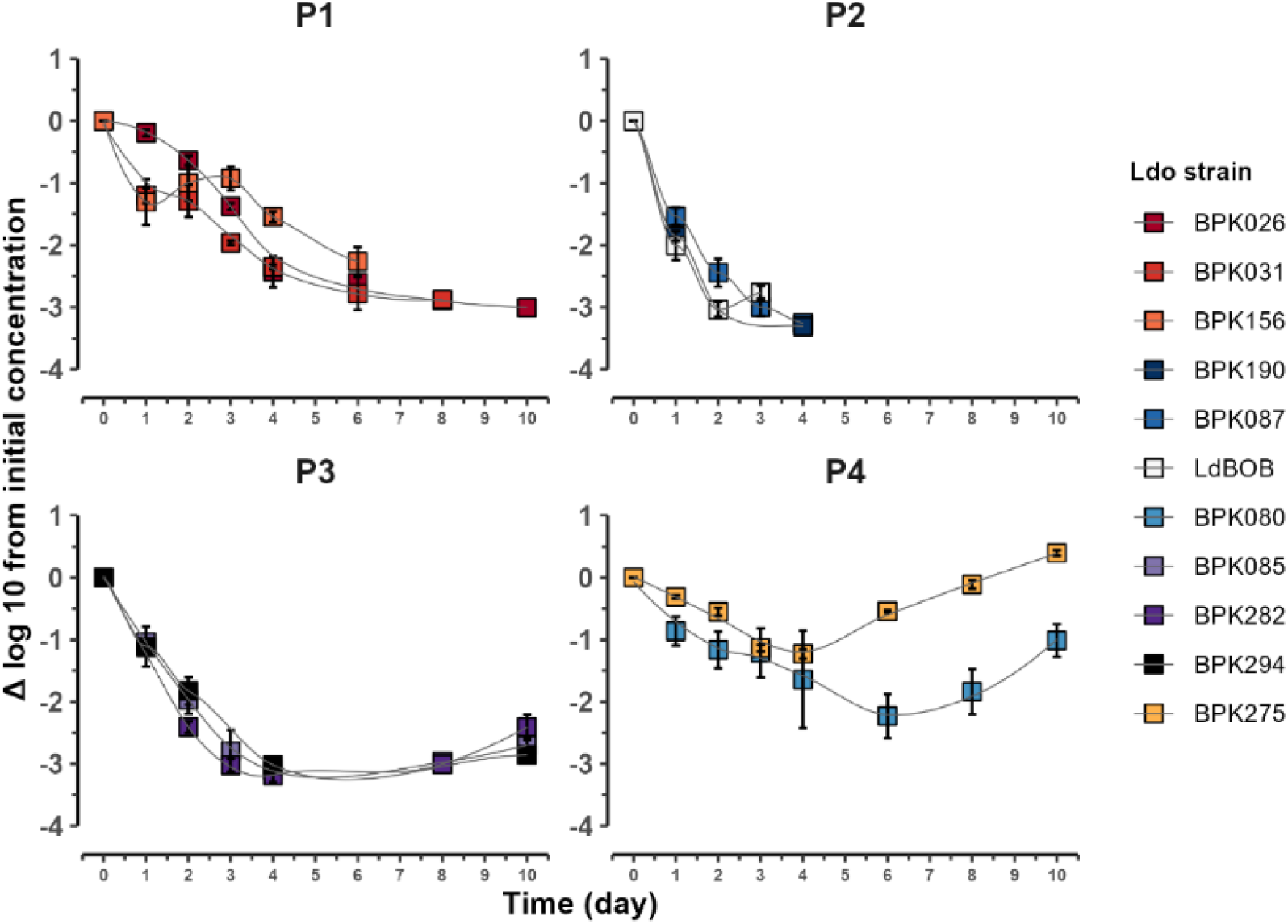
Patterns of survival of *L. donovani* strains under normally lethal PAT concentrations. Time-kill curves under prolonged PAT exposure (see separate strain-specific curves in Supp. Fig.2) showed four different viability profiles (Profile 1-4). Data represents the reduction in promastigote concentration relative to the initial inoculum, the viable parasites were detected by flow cytometry. The results reflect the mean ± SEM of three biological replicates.

Profile 1 (P1) observed in the ISC1 parasites (BPK026, BPK031, BPK156) was characterized by continuous reduction of viability throughout the whole period of exposure to PAT, with the majority of the parasites killed in the first 6 days, while a small remaining population (in the range of 0.7-3.4%) was killed at a slower rate over the last 4 days, allowing those cells to persist until the end of exposure. In profile 2, P2, (BPK190, BPK087, and LdBob) we observed rapid killing of almost the entire population of cells in the first 2 days (1.8-11.0% survived beyond day 2), followed by a slower killing over the following days, with no viable parasites found by day 8. Profile 3 (P3) exhibited by BPK085, BPK282, and BPK294 could be distinguished by fast reduction in viability over the first 3-4 days reaching viability 1.9-3.4%, followed by a slower killing rate until days 6-8. Surprisingly, in the last 2 days of exposure to PAT, a clear increase in the number of viable parasites was observed reaching 5.5-14.9% at day 10. Finally, the last profile (profile 4, P4) displayed by BPK080 and BPK275 is defined by mild reduction in viability, followed by a short period of no visible growth, then a subsequent increase in the number of viable cells. BPK275, a strain with the two genomic pre-adaptations to antimony was the least affected by PAT having a viability of 70% at its lowest value (Supp. Table 2).

### *L. donovani* lines adopt a quiescent state to survive PAT pressure

Given the apparent reduction in growth in all tested lines, we addressed whether the PAT-exposed cells adopt a quiescent state as a survival mechanism to prolonged drug exposure. We addressed this first by analyzing the reduction of rEGFP expression -a proxy of metabolic activity-^36^ in all the lines exposed to PAT at day 2, and 4 (PAT D2 and PAT D4, respectively), as compared to proliferative promastigotes. We included, as a positive control, stationary promastigotes (No PAT D4), which are known to have properties of quiescent cells ^24^. In all lines of P1/P4 we observed a progressive reduction of rEGFP under PAT pressure, reaching comparable or slightly lower levels to those observed in No PAT D4 (Fig. 2A). In parasites belonging to P2, and P3 measurements were possible only at D2 (all lines in P3 and only BPK087 in P2) due to rapid killing in the lines of these profiles, which resulted in an insufficient number of viable cells for reliable quantification of rEGFP expression. As in other lines, we detected reduction in rEGFP expression at D2 in all three P3 lines. Surprisingly, we observed an upregulation of rEGFP in BPK087 suggesting an inability to enter quiescence under PAT pressure (62.7% of increase, Fig.2A). As expected, all lines that were able to survive drug pressure, resumed normal growth after transfer to standard growth conditions without PAT (Supp. Table 3).

**Figure 2.**
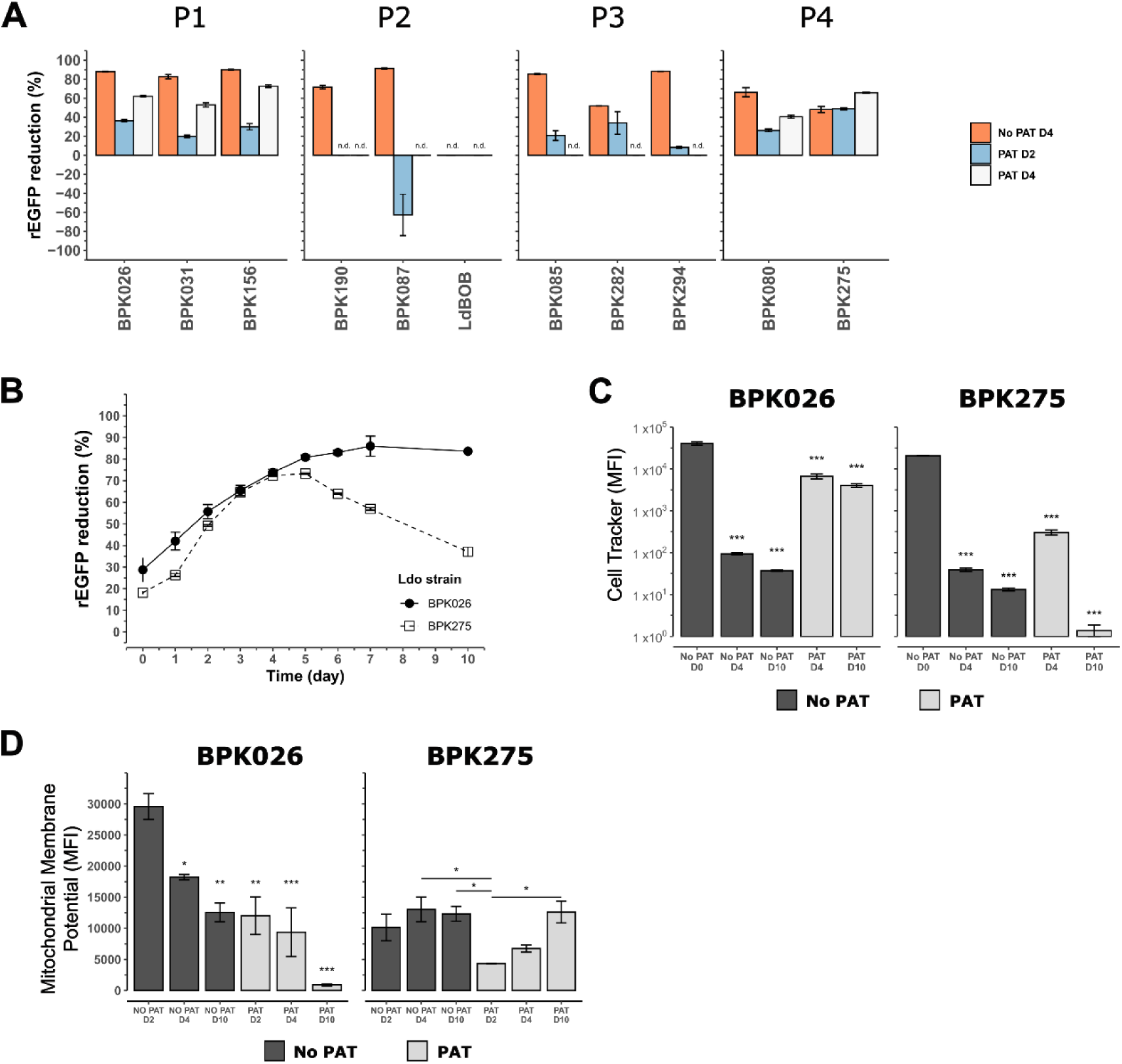
Quiescence features of *L. donovani* promastigotes under prolonged PAT exposure. (**A**) rEGFP expression reduction in percentage in P1-P4 profiles under normally lethal doses of PAT exposure for 2 (PAT D2) and 4 (PAT D4) days. A group without exposure for 4 days was included (No PAT D4). In certain samples, rEGFP expression data is not reported due to a limited number of viable cells, indicated as ‘n.d.’ (not determined). (**B**) rEGFP expression reduction in percentage in BPK026 and BPK25 from day 0 to 10. (**C**) Proliferation status measured by CellTracker in promastigotes from BPK026 and BPK275 under PAT exposure (PAT) with the respective no-drug controls (No PAT). The results were compared to the initial staining point of the parasites (No PAT D0). (**D**) Measurement of the mitochondrial membrane potential with MitoProbe ™ DilC by flow cytometry in promastigotes exposed for 2, 4, and 10 days to strain-specific PAT normally lethal dose (PAT); and the respective controls without drug (No PAT). The results were compared to promastigotes at the logarithmic state (No PAT D2). (**A,B**) The rEGFP expression reduction in percentage was calculated by comparison of the signal of each group to promastigotes without PAT exposure at the logarithmic state. The data reflects the mean ± SEM of three biological replicates. The asterisks represent the statistically significant difference after Tukey’s post-hoc test; * P < 0.05, ** P < 0.01, *** P < 0.001.

To confirm the quiescent status of the viable cells under PAT, we selected two lines, BPK026 and BPK275, which present the most extreme viability profiles (P1, and P4, respectively) and extreme genomic pre-adaptations to PAT (no known genomic pre-adaptations, and H-locus amplification together with the AQP1 indel associated with antimony resistance, respectively) for more in-depth characterization. First, we measured daily rEGFP expression until day 7, and at the last day of PAT exposure. Both lines showed an overall rEGFP decrease among viable cells over the first 4 days of PAT pressure (Fig. 2B), matching the observed growth arrest (Supp. Fig. 4A). This decrease continued for BPK026 over the next 6 days, reaching a value of 83.1% of reduction in expression. In contrast, rEGFP expression for BPK275 remained stable until day 5, followed by a clear increase on day 10 (Fig. 2B). This is consistent with the apparent reactivation and increase in the percentage of viable and proliferative cells towards the end of drug exposure (Supp. Fig. 4B). Second, we assessed whether parasite replication is arrested/reduced using CellTracker (tracking cell proliferation) combined with SYTOX Blue dye (viability, Fig. 2C). Each round of replication dilutes the CellTracker dye resulting in decreased fluorescence signal, thus fluorescence levels are expected to remain unchanged in non-proliferating cells. In this assay, logarithmic promastigotes were labelled by CellTracker (D0) and either exposed to PAT at a normally lethal dose (PAT) or grown in the control medium (No PAT), and the fluorescence signal was evaluated during short-(D4) and long-term (D10) PAT exposure. In both lines, No PAT parasites showed clear proliferation as indicated by a decrease in fluorescence, reaching 1.97 - 1.57 to 1.59 - 1.12 log_10_ values for BPK026 and BPK275, respectively (Fig. 2C). PAT-exposed parasites by contrast showed less dilution (Fig. 2C). In BPK026, the CellTracker signal decreased slightly from 4.61 to 3.82 log_10_ values at PAT D4, and remained almost the same at PAT D10 (3.61 log_10_ values) indicating an overall low/no proliferation of PAT-exposed cells. In the case of BPK275, the signal in PAT D4 (2.48 log_10_ values) decreased as compared to D0 (4.31 log_10_ values), but it was still higher than fluorescence in the No PAT D4 parasites (1.59 log_10_ values). No CellTracker signal was detected in PAT D10; this indicates that during the early exposure to PAT, BPK275 parasites transiently reduce their proliferation, while resuming active proliferation later.

Next, we investigated the mitochondrial membrane potential as an indicator of mitochondrial activity, which is known to be reduced in quiescent *Leishmania* cells ^24^. We labelled promastigotes at different time points under PAT exposure and their respective controls, with MitoProbe™ DilC, a permeable cationic dye that accumulates across the mitochondrial membrane when the interior of the mitochondrion is negatively charged compared to its outside (Fig. 2D). In the absence of PAT, BPK026 showed high potential in logarithmic promastigotes, and this decreased significantly in the stationary phase. Unexpectedly, in BPK275, the mitochondrial membrane potential was much lower in logarithmic parasites in comparison to BPK026 (29,569 ± 2,071 and 10,142 ± 2147 MFI values, in BPK026 and BPK275, respectively, Fig. 2D). Under PAT pressure, patterns were also markedly different between lines. In BPK026, early PAT-exposed promastigotes showed a similar potential to late stationary (D10) promastigotes, which further decreased to 899 ± 165 MFI values at day 10 (Fig. 2D). In the PAT-exposed BPK275, the mitochondrial potential was the lowest at D2 (4,326 ± 47 MFI value), and it gradually increased reaching 12,611 ± 1,741 MFI value at D10 (Fig. 2D).

Thus, based on the rEGFP, CellTracker, and mitochondrial potential measurements, we concluded that both lines exhibited features of quiescence during their exposure to normally lethal doses of PAT. Though in BPK275, the line with genetic resistance to PAT, the period of quiescence was transient, and the quiescence state appeared to be less deep. The accumulated data demonstrates that although different dynamics of killing could be observed in *L. donovani* promastigotes exposed to lethal doses of PAT, in all lines that survived PAT pressure, adoption of a quiescent state was associated with survival.

### The impact of transition through quiescence during exposure to PAT on antimony susceptibility depends on the presence of genomic pre-adaptations to antimony in analysed *L. donovani* lines

Although the quiescent state was found in all but one of the studied lines, the different killing profiles observed earlier, suggested different survival mechanisms. Specifically, P1 parasites seemed to behave similarly to classical bacterial persisters with no growth during exposure to the drug, while P3 and P4 parasites were able to resume growth under drug exposure after a short transition through the quiescent state, suggesting that a small subpopulation of these resistant parasites became hyper-resistant. To ascertain whether drug-pressure was selecting for altered genome sequences of parasites isolated post-exposure to PAT, we analysed the PAT-recovered parasites with particular focus on their PAT susceptibility and potential genomic alterations.

First, we measured the PAT susceptibility of the cells that recovered from the PAT exposure by comparing the IC_50_ without drug exposure (No PAT) and after removal of PAT and recovery of growth (Post-PAT) (Table 2). As reported previously, we found that the P1 parasites (ISC1 group) showed the highest susceptibility, with IC_50_ values ranging between 9 ± 0.6 and 19 ± 1.2µM. The P3 and P4 parasites exhibited a lower susceptibility, with IC_50_ values ranging between 46.4 ± 0.5 µM and 137.1 ± 11.9 µM. When control and Post-PAT conditions were compared, two patterns were observed: (i) P1 lines did not show changes in IC_50_ after the exposure to PAT; (ii) in contrast, the P3 and P4 lines showed a significant increase of IC_50_ in Post-PAT conditions. The result observed for the P1 parasites is consistent with the classical behaviour reported for persister cells in bacteria. However, the extreme increase in IC_50_ values in such a short period of time observed for P3/P4 parasites was unexpected, but consistent with the ability of these parasites to grow in the presence of PAT.

**Table 2.**
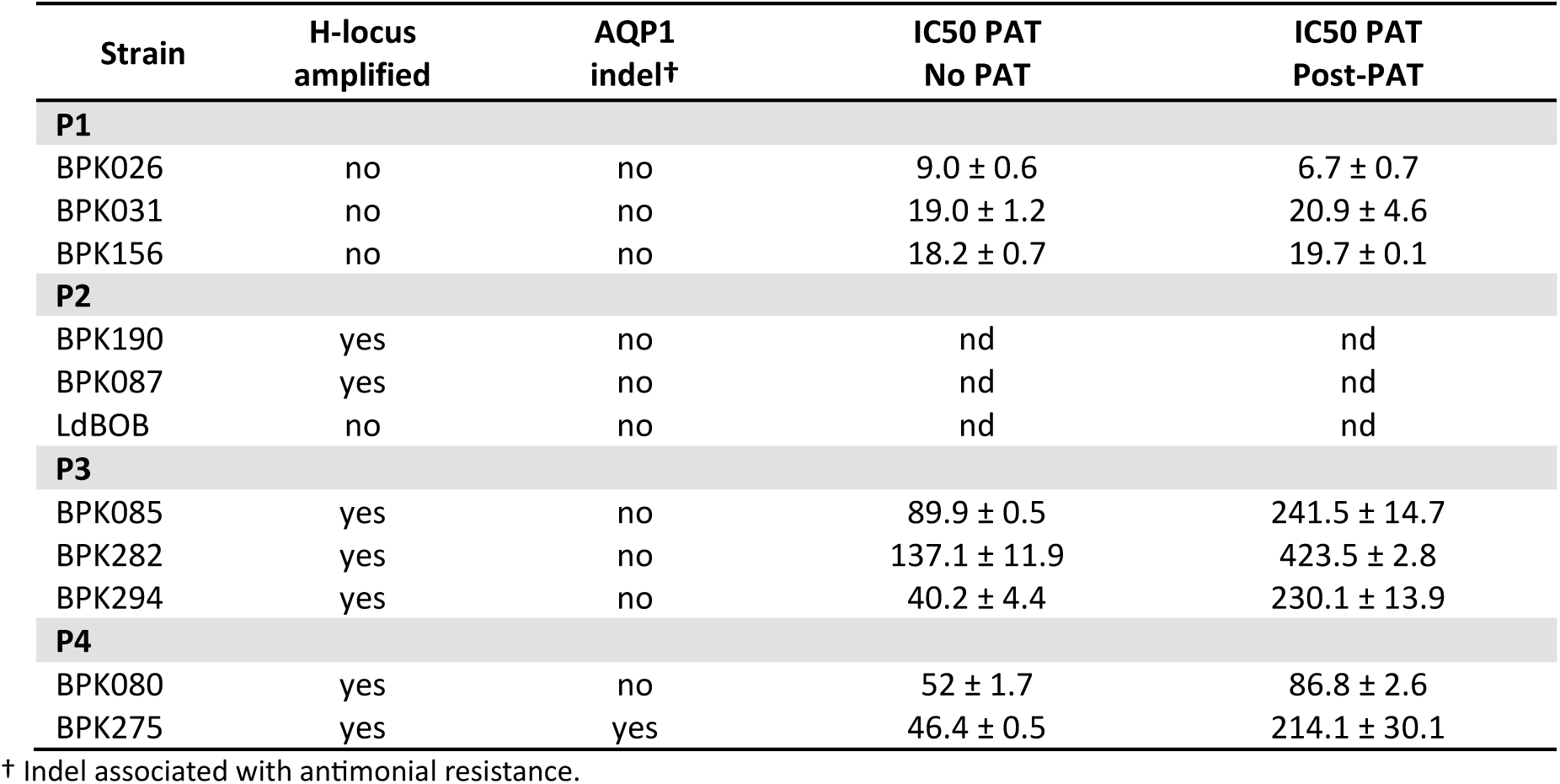
Drug susceptibility of *L. donovani* promastigotes (IC_50_, µM) of ten ISC and one reference strain measured by resazurin assay after removing normally lethal dose 10-day of PAT (post-PAT); the control (No PAT) was maintained in parallel and not exposed to the drug. For three lines that did not recover after drug exposure, the data is not reported and denoted as ‘n.d.’ (not determined). The data is presented as the mean ± SEM of three biological and technical replicates.

We then, we sequenced and compared the genome of control and Post-PAT, in triplicates, in order to search for possible signatures of drug resistance. In our analysis, we focused on potential changes of SNPs, INDELs, copy number of individual genes and the H-locus and somy values. Because of ISC1 strains/P1 parasites having much higher number of heterozygous SNPs in comparison to other strains ^16^, we carried out the analysis of P1 and P3/4 parasites separately. In both P1 and P3/4 lines, we did not observe consistent (*i.e.* present in all three replicates) SNP/indel changes (Fig. 3A,B; Supp. Fig. 5) changes. Similarly, there were no consistent changes in the depth of individual coding sequences (Fig. 4A,B; Supp. Fig. 6), including the MRPA genes (Supp. Table 4) when No- and Post-PAT parasites were compared. However, when we analyzed somy changes in Post-PAT parasites as compared to non-exposed lines, we observed a general decrease of somy of several chromosomes in both P1 and P3/4 lines (Fig. 5; Supp. Fig. 7). Focusing on the somy changes that we consider biologically significant (S values difference of at least 0.5, p-value≤0.05), we detected: i) decrease in at least one chromosome in all P1 parasites, namely chr 33 and chr07 in BPK026, chr 23 in BPK031 and chr 09 in BPK156. In addition, five chromosomes, chr 02, chr 08, chr 14, chr 32and chr 35 were decreased in BPK085, and chr 26 in BPK294 ii) no significant increases in chromosome copy number were found in P1 parasites, while we observed a significant increase in copy number of chr 6 and ch31 in BPK282 and BPK275. In addition, an increase (albeit not reaching statistical significance) in chr 31 was found in BPK294, where chr 07 was also increased. No significant aneuploidy changes were detected in BPK080.

**Figure 3.**
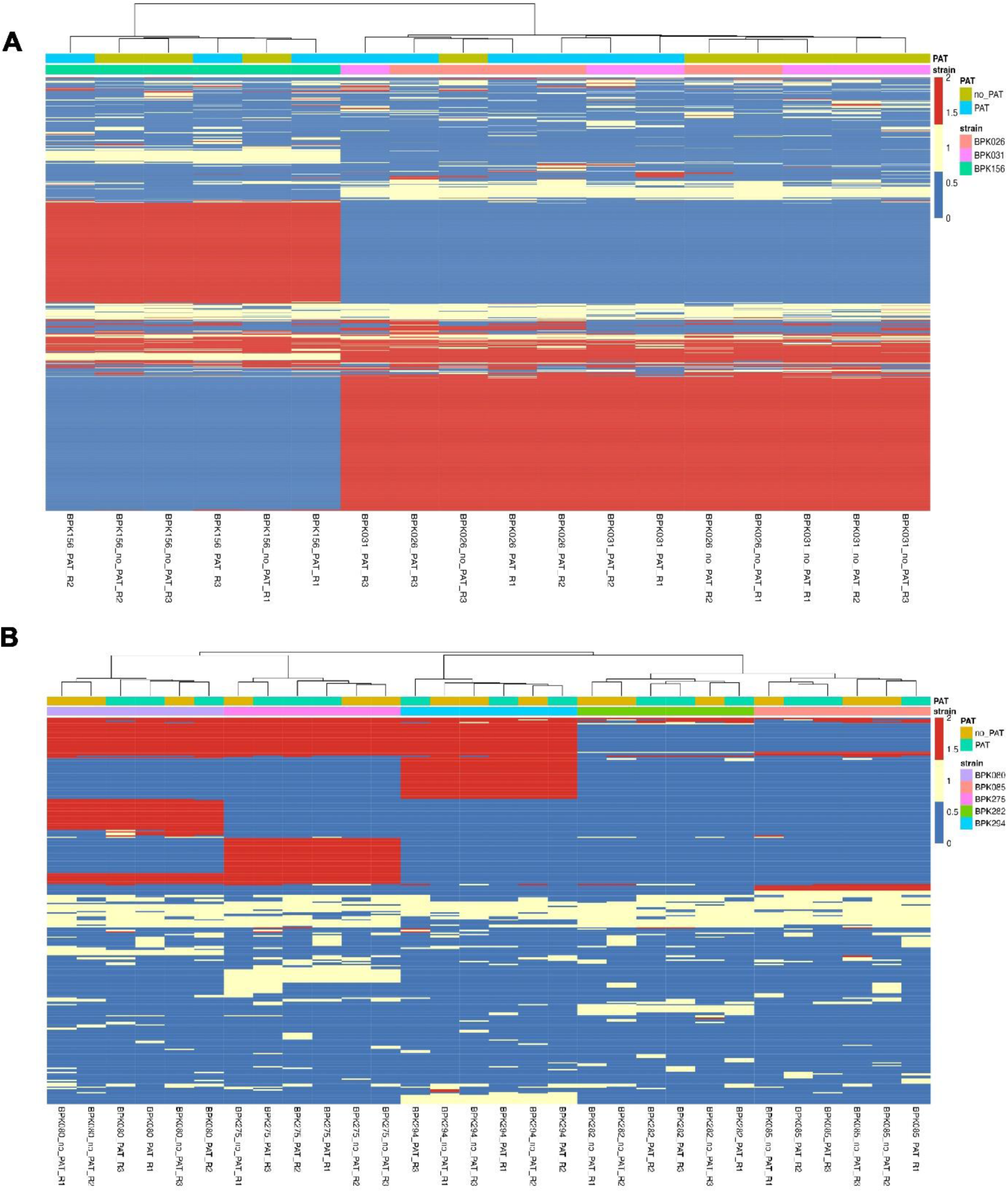
Heatmap illustrating the distribution of single nucleotide polymorphisms (SNPs) across strains belonging to. (**A**) Profile 1 and (**B**) Profiles 3/4. SNPs that exhibited no variation across the 30 Profiles 3/4 samples or the 18 Profile 1 samples were filtered out. The color scheme categorizes SNPs as follows: blue represents the absence of a SNPs, yellow indicates heterozygous SNPs, and red denotes homozygous SNPs. Hierarchical clustering of samples (columns) and SNPs (rows) was performed using the complete linkage method with Euclidean distance. A color-coded bar above the heatmap indicates whether each strain was exposed to PAT, alongside the strain names. The data represents three biological replicates (R1, R2, R3) after removing normally lethal doses of PAT (PAT), and their respective control without drug (no_PAT).

**Figure 4.**
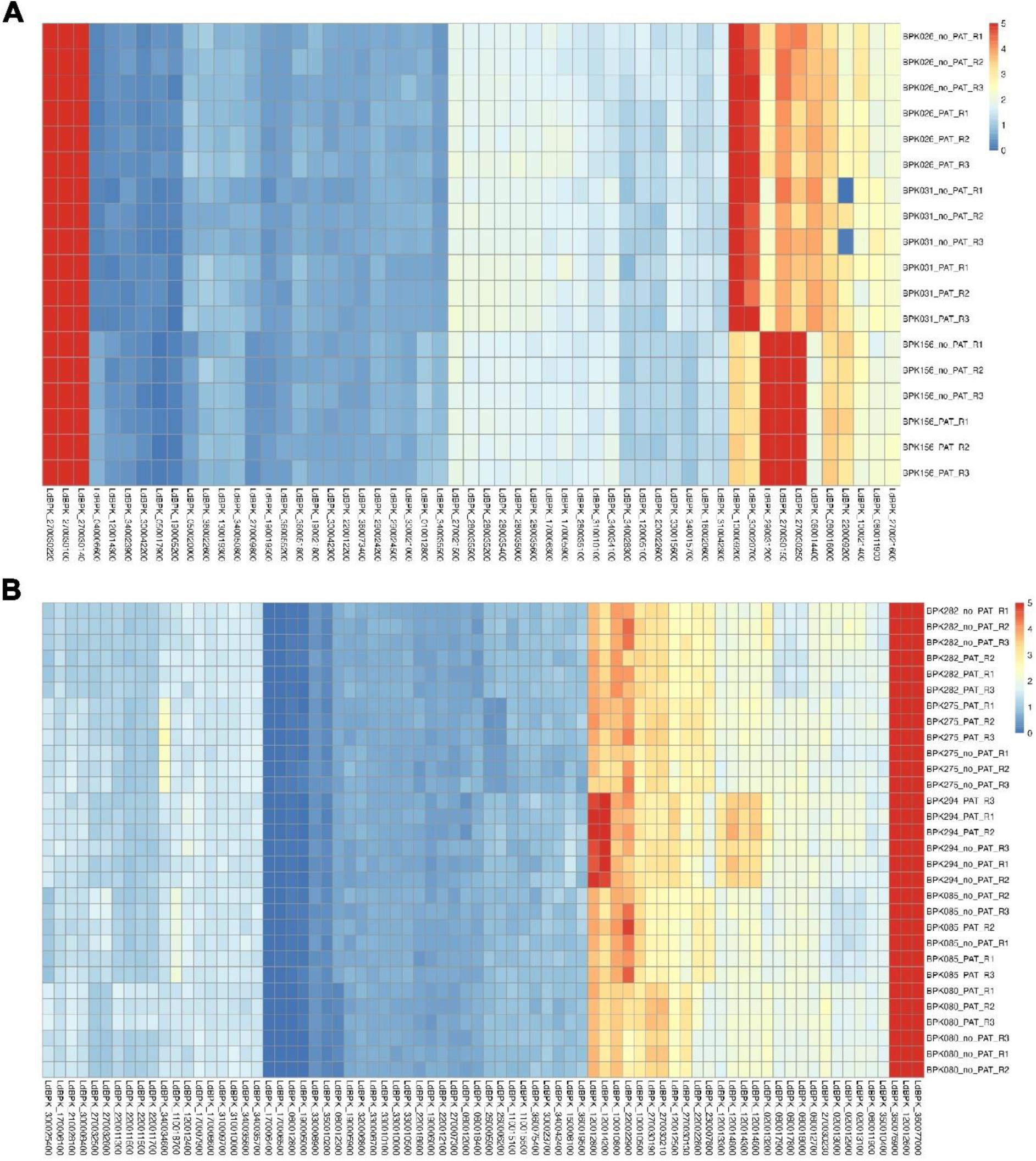
**Heatmap of the** (**A**) Profile 1 and (**B**) Profiles 3/4 strains, displaying CNV values for genes with a statistically significant 50% increase or decrease in CNV. Statistical significance was determined using a Student’s t-test to compare CNV values between PAT-exposed (PAT), and non-exposed replicates (No_PAT) for each strain. Three biological replicates were included (R1,R2,R3) for each strain and drug exposure.

**Figure 5.**
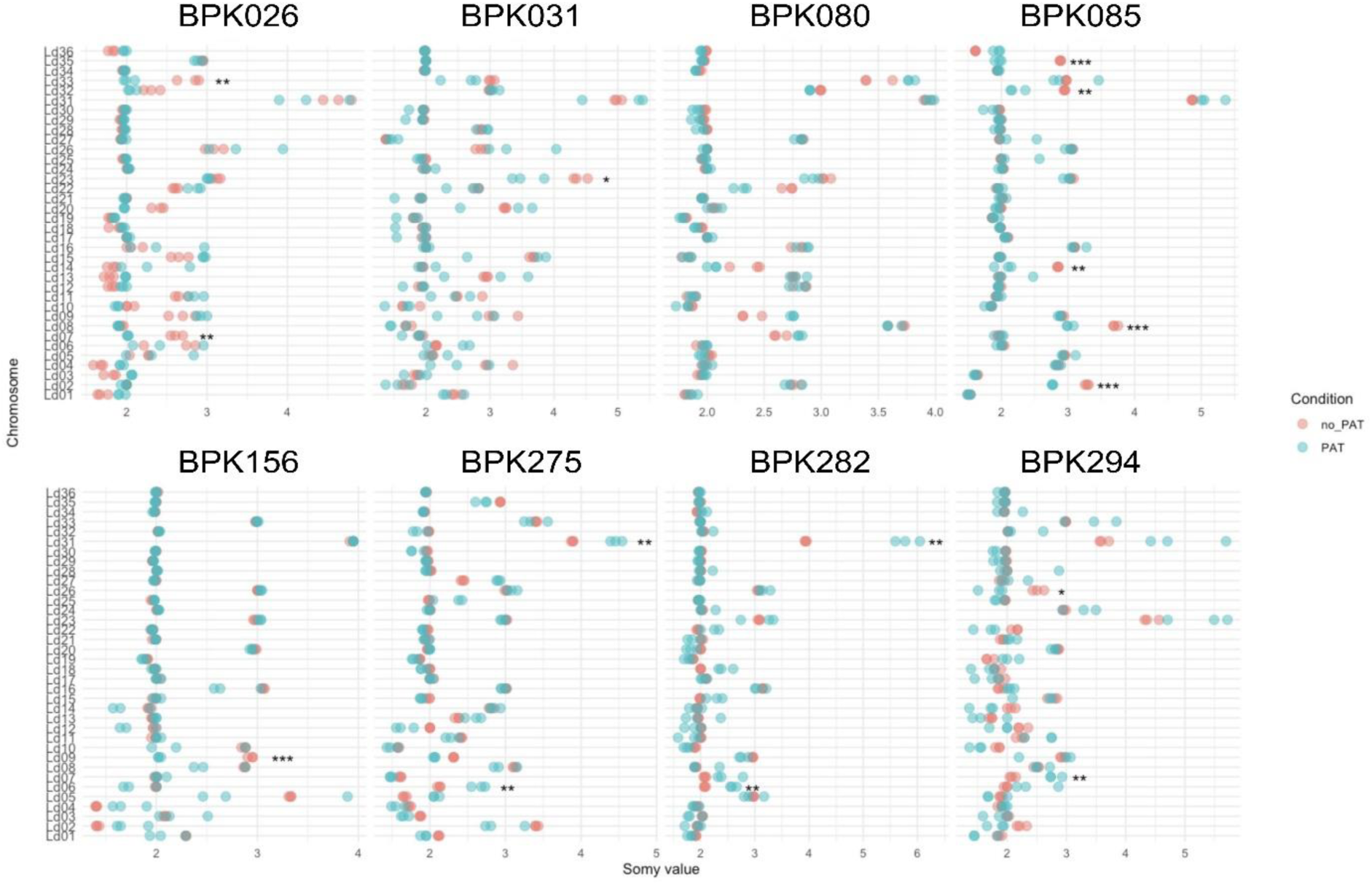
Dot plot containing the somy values per chromosome and per strain. For each chromosome and each strain, the somy value is indicated in the non-exposed samples (red, No_PAT) and the PAT exposed sample (green, PAT). Statistically significant somy differences using student t-test between non-exposed versus PAT-exposed samples are indicated by asterisks per chromosome and strain (* < 0.05, ** < 0.01, *** < 0.001). The data represents the values of three biological replicates.

In summary, no consistent genomic alterations could be detected in either P1 or P3/4 parasites. This further supports persister-like nature of a small population in P1 parasites that survive normally lethal doses of PAT. In the case of P3/P4 parasites, we report here detection of a unique form of transient persisters with the ability to retain their reduced PAT susceptibility in drug-free conditions showing no discernible genetic change when compared to the precursors. Further investigation, possibly into epigenetic modification is required to ascertain the mechanism allowing the growth in PAT and increase of IC_50_ in recovered P3/P4 parasites.

Overall, the impact of the transition through quiescence, on drug susceptibility differed, depending on the presence of genomic pre-adaptations to antimony. Lines without any genomic pre-adaptation exhibited features of a classical quiescence phenomenon linked to the presence of persisters, where increased tolerance to a drug is restricted to the quiescence phase. Consistent with this notion, no clear drug resistance signatures were detected in Post-PAT parasites. While in the lines that were genomically pre-adapted, after a short period of quiescence, a subpopulation of more tolerant cells was able to resume growth even in the presence of PAT. These emergent cells had a level of PAT resistance beyond the level of the parent clone in which quiescence was initially induced. However, no consistent genomic changes could be detected in the analyzed P3/P4 parasites. This suggests a presence of a unique form of adaptation in these lines, in which the exact molecular mechanism underlying the reduced PAT susceptibility remains to be characterized.

### PAT-tolerant promastigotes show cross-tolerance to other drugs

In bacteria, entering into a non-proliferative physiological state after the exposure to one drug, can provide tolerance to other antibiotics ^6–8^. Hence, we tested whether the observed PAT tolerance, associated with the acquired quiescent state in the BPK026 and BPK275 lines, would also provide tolerance to other known antileishmanial compounds. We, therefore, evaluated the susceptibility of PAT-tolerant promastigotes of both BPK026 and BPK275 towards 5 compounds with different modes of action: amphotericin B (AMB), miltefosine (MIL), bortezomib (BOR), carbonycil (FCCP) and paromomycin (PARO). Briefly, i) AMB binds ergosterols and potentially destabilises membranes, ii) MIL is a lipid binding compound and likely also acts through destabilization of membranes, iii) BOR is a proteasome inhibitor, iv) FCCP is a protonophore, which can permeabilise the mitochondrial membrane, and v) PARO is a probable protein synthesis inhibitor ^38–42^. First, we found that the IC_50_ values for each of these compounds were comparable in BPK026 and BPK275 (Supp. Table 5). To assess the protective effect of PAT-associated tolerance, we first triggered quiescence by exposing the promastigotes to PAT (strain-specific normally lethal doses) for 5 days, followed by 5 days of simultaneous exposure of PAT and the additional drug (serial exposure (S), Supp. Table 6). Secondly, we added both PAT and the second drug from the very beginning (combination exposure (Co), Supp. Table 6). The drugs were applied at their respective IC_50_ concentrations in both Co and S conditions. As a control exposure, we repeatedly treated with PAT during the first and second exposure (Supp. Table 6). To assess the killing effect of these drugs, we quantified the % of viable promastigotes in relation to the control exposure (C).

In the case of BPK026, two main patterns were observed. On one hand, with AMB, MIL, and BOR, there was a strong reduction of the viability ratio in respective combination exposures (0.001 to 0.34 range), while in the serial exposures, the viability was comparable to the levels in control treatment (0.79-0.82 values in S, and 1.0 in C; Fig. 6A). This indicates that PAT-triggered BPK026 quiescent cells gain tolerance to AMB, MIL, and BOR. On the other hand, with FCCP and PARO, viability remained similar to control exposure in both Co and S conditions, suggesting that used concentrations of FCPP and PARO were too low to enhance PAT killing (Fig. 6A). Results were confirmed by the reversibility assay: *i.e.* the combination of PAT+AMB for which growth was observed in only one out of three replicates, while all three replicates demonstrated proliferation in the case of serial exposure (Table 3). In the case of BPK275, results were different and 5 patterns were observed (Fig. 6A): (i) for BOR, a low viability ratio in Co (0.047 ± 0.02 value), and comparable viability in the C exposure and S exposures, suggesting cross-tolerance to BOR in PAT-exposed quiescent cells, ii) lower viability ratios for AMB (both Co and S exposures) in comparison to C, but the viability in S was significantly higher than in Co, suggesting some level of protection against AMB provided by quiescence of PAT-exposed cells (iii) same viability as control for MIL (both Co and S conditions), indicating that used MIL concentrations were too low to enhance PAT killing, (iv) for FCCP, although low viability in S was observed, microscopic examination revealed an abundance of parasites, suggesting that parasites likely died due to nutrient deprivation, and (v) for PARO a higher viability ratio in Co exposure in comparison to C and a ratio close to the control under serial exposure, suggesting that PARO and PAT act synergistically to trigger the presence of quiescent cells. In all conditions, the reversibility assay revealed proliferation in the 3 replicates (Table 3). Collectively, these results show that PAT-tolerant cells also show tolerance to other drugs, but the degree of this cross-tolerance is drug- and strain-dependent.

**Figure 6.**
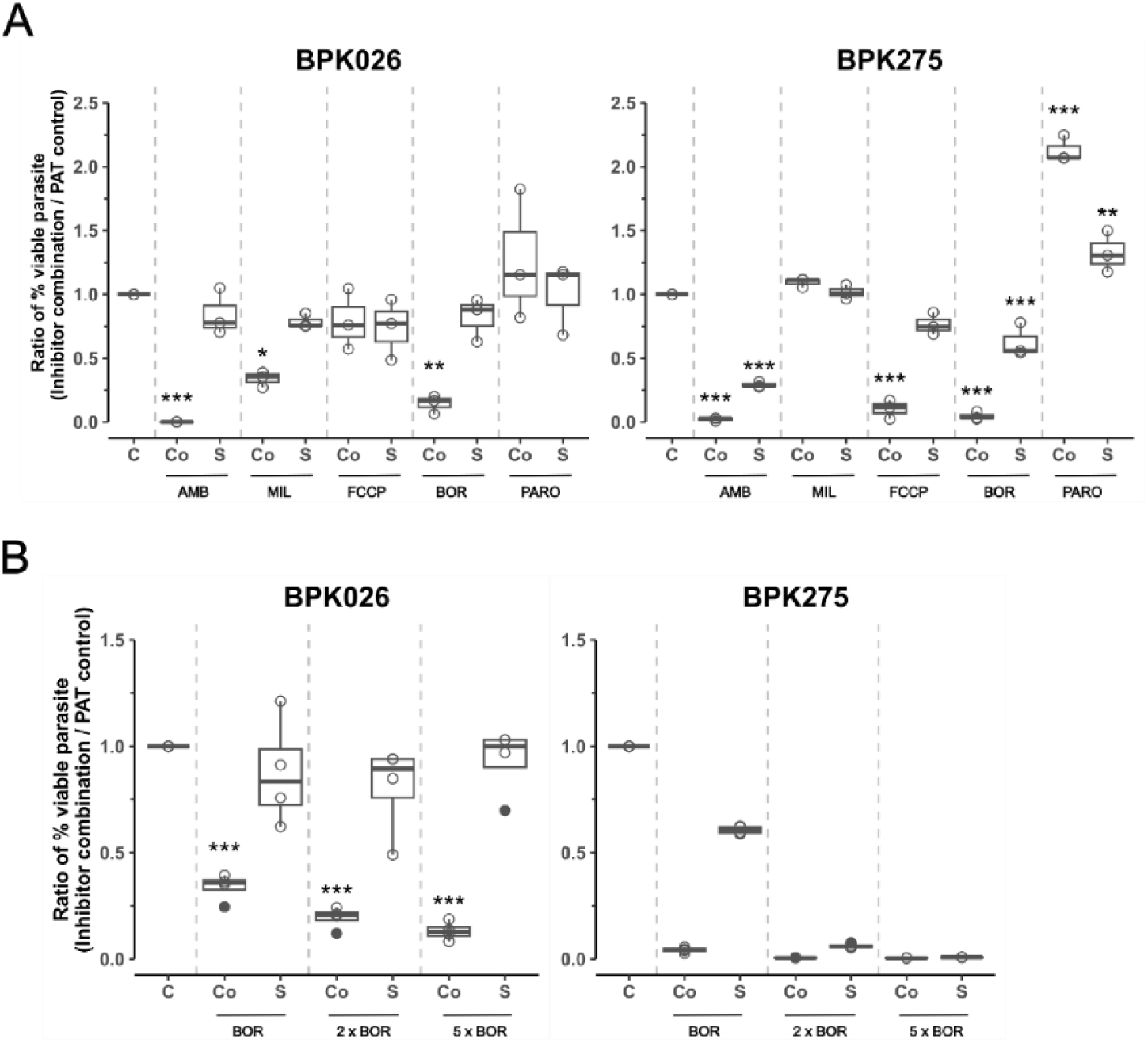
Survival of PAT-tolerant *Leishmania* promastigotes when exposed to other drugs. (**A**) Ratio of percentages of viable parasites submitted to different compounds (IC50) vs normally lethal doses of PAT (control, C) and following two drugging schemes (Co, combination and S, serial); (i) MIL, Miltefosine; AmB, Amphotericin B, PARO, Paromomycin; BOR, Bortezomib and FCCP, Carbonycil. Plotted data shows the mean ± SEM of three biological replicates for each combination or serial. (**B**) Ratio of percentages of viable parasites submitted to BOR (1x IC50, 2x and 5x IC50) vs normally lethal doses of PAT (control, C) and following two drugging schemes (Co and S). Plotted data shows the mean ± SEM of four biological replicates for each combination or serial. The asterisks indicate statistically significant differences for each scheme compared to the control, analyzed by Tukey’s post-hoc test (**A**) and Student’s t-test, (**B**)* P < 0.05, ** P < 0.01, *** P < 0.001.

**Table 3.**
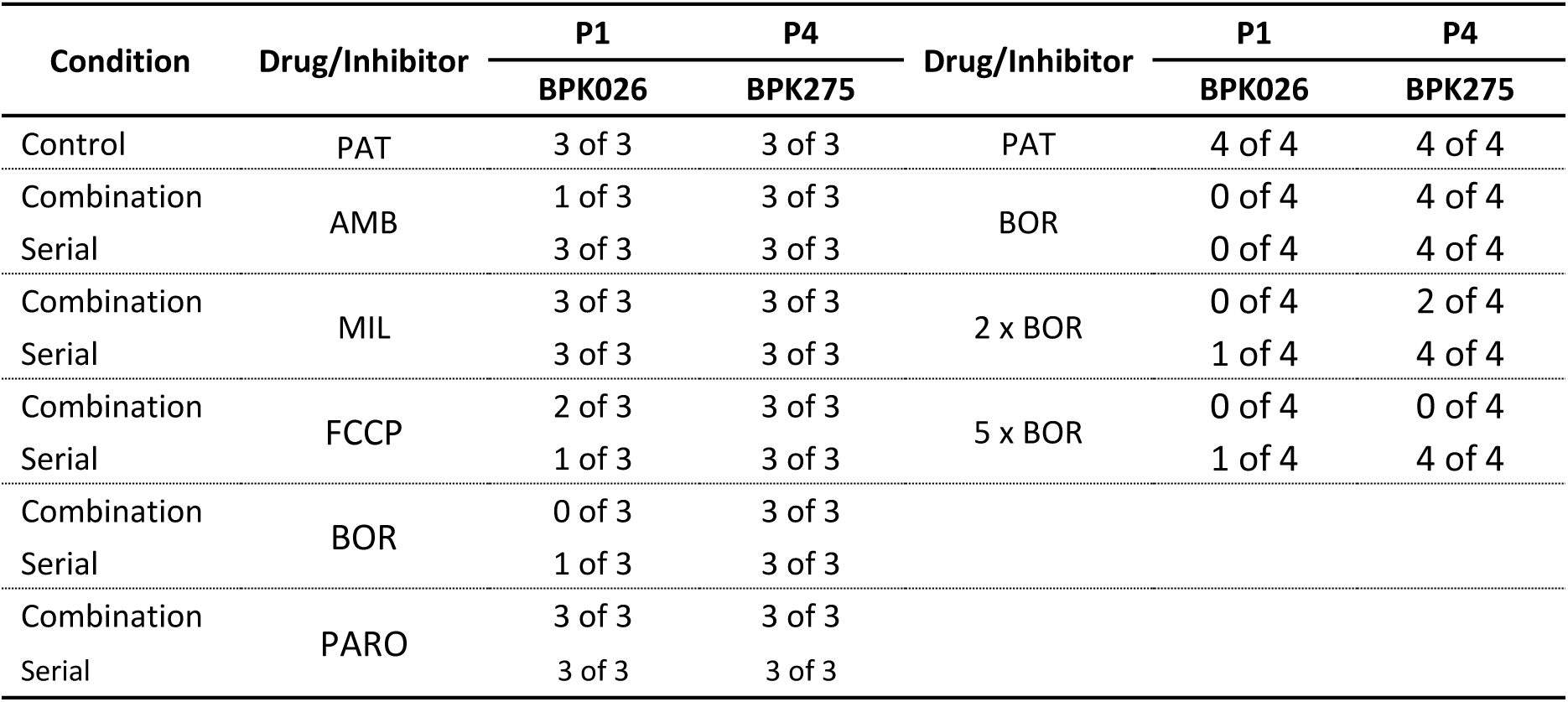
Reactivation of parasite proliferation (numbers out of 3 or 4 replicates) after exposure to a combination of PAT and another drug or serial drugging (see experimental design in Supp. Table 6 or Supp Table 7).

To analyze the cross-tolerance further we focused on protection against BOR, for which we also executed combination and serial exposure and compared the effects of applying BOR at 3 different concentrations: at 1x, 2x, and 5x IC_50_ of BOR (see experimental design in Supp. Table 7). In the case of BPK026, viability ratios slightly decreased with the increasing concentration of BOR in the combination, condition, but this did not affect the observed cross-tolerance in S exposure (Fig. 6B). In contrast, in BPK275, viability ratios decreased with the increase of BOR concentration in both conditions, with almost no viable cells detected in the two highest concentrations of BOR. (Fig. 6B). The reversibility assays showed that none of the Co replicates could grow, while a few replicates in the S condition could re-proliferate in BPK026. By contrast, for BPK275, all replicates in the S condition could reactivate, while only two or none were able to proliferate in the higher concentrations of BOR in Co exposure (Table 3). This indicates that cross-tolerance to BOR could be observed for both strains, but it was more pronounced for the antimony resistant BPK275 line.

## Discussion

This study demonstrates that genetically different *L. donovani* strains use quiescence as a survival strategy under normally lethal doses of trivalent antimony. In most lines, upon exposure to PAT a small non-replicating population with reduced metabolism was observed and had the ability to survive the treatment. A quiescence state was also found in an antimony resistant *Leishmania* line, in which a majority of cells survived drug exposure. Quiescence was detected in all parasites with genomic pre-adaptation to antimony, and these cells could also resume growth while still under treatment exposure, enhancing the level of resistance to PAT in so doing. Quiescence, therefore, appears to be a widespread phenomenon in *Leishmania* parasites exposed to normally lethal doses of a drug.

Here we triggered entry into quiescence by exposure to antimony, analogous to triggered persistence in bacteria rather than spontaneous persistence ^4^. All lines surviving normally lethal doses of PAT, showed a bi-phasic time-kill curves typical for bacterial persister cells ^43,44^, though in two lines, BPK275 and BPK080, the subpopulation of PAT tolerant cells was larger than typical. Detailed analysis of the lines with the most extreme survival phenotypes revealed several properties typical of persisters, including reduction of metabolism, proliferation and mitochondrial membrane potential with no apparent genomic changes and cross-tolerance to other antileishmanial drugs. However, the ability to resume growth after a transient adoption of quiescence observed in lines with pre-existing antimony resistance is inconsistent with classical definition of bacterial persisters, since these lines acquire a further reduction in sensitivity to drug during the passage through quiescence. Because of this unique mixture of observed phenotypes in the quiescent *L. donovani* lines, we collectively call them persister-like cells.

Parasites without known antimony resistance, when triggered to enter quiescence over a 10-day period remain quiescent while still under drug treatment, and resume growth only when transferred to antimony-free medium. By contrast, *Leishmania* that had harboured genetic mutations that yielded resistance to antimony, also entered quiescence when exposed to antinomy at a dose beyond that to which they are normally resistant. However, these cells were able to re-enter growth phase whilst in medium containing drug. Remarkably, these parasites that resume growth whilst still in medium containing antimony are actually more resistant to the drug than they were before having passed through quiescence. Although we do not have a molecular explanation for this, it is possible that the MRPA pump which removes antimony from the cells ^45^, and is amplified as part of the H-locus in resistant cells is expressed at still higher levels independently of gene copy number (which was not further amplified during the quiescence period). Given that drug-efflux is a common mechanism of resistance in these strains, it is possible that the resistant lines reach a point at which the intracellular levels of antimony decrease below a threshold required to retain the triggered quiescence phenotype. It will be interesting to explore this in future work.

We also tested the ability of cells in which quiescence was triggered by PAT to tolerate exposure to a range of other drugs. By adding 5 other compounds, with different modes of action, at their IC_50_ levels, alongside PAT on day zero, we were assessing their toxic effect on non-quiescent cells. By contrast, by adding the secondary drugs five days after the addition of PAT, we were testing their ability to kill pre-selected quiescent cells. AMB, MIL and BOR had significantly reduced ability to clear cultures if added after 5 days while this was not the case for PARO and FCCP. Mechanistically we had assumed AMB might not have different impact on quiescent cells as it permeabilised membranes after binding ergosterol. However, quiescent cells appear to be less sensitive to the drug, possibly indicating membrane remodelling during quiescence, which might also explain why quiescent cells also showed tolerance to MIL. In the case of the proteasome inhibitor BOR, the general decrease in protein turnover in quiescent cells may explain why cross-tolerance is seen to this compound. FCCP interferes with proton-dependent transport processes at the mitochondrial and plasma membranes and given the requirement to sustain key metabolic processes the fact that quiescent cells retain sensitivity was not a surprise. As an inhibitor of protein synthesis paromomycin likely continues to act on processes crucial to the viability of quiescent cells, which would explain the absence of cross-tolerance to this drug.

Cross-tolerance to several antifungal compounds has been observed in pathogenic fungi and was linked to changes in aneuploidy patterns and not the persister state ^46,47^. Interestingly, it has been reported by us and others that aneuploidy is very dynamic and can be adaptive in *Leishmania* ^19,32,33,48,49^. We analyzed genomic changes in both lines that showed cross-tolerance, but only in the recovered parasites grown in drug-free medium, where no consistent chromosome copy number variations were found. However, given the aneuploidy dynamics and mosaicism in *Leishmania* we cannot exclude that some chromosome copy number changes contributed to the detected cross-tolerance in persister-like cells.

In our experimental set up, the impact of transition through quiescence extends beyond the duration of the non-proliferative period. We found that the lines with pre-existing adaptations to antimonials (through amplification of a drug exporter and/or inactivation of AQP1 transporter) emerged with reduced PAT susceptibility after recovering growth in drug-free condition. Given the lack of detected genetic changes in those parasites, and the isogenic nature of the used lines we propose that a form of resetting mechanism takes place, perhaps through epigenetic reprogramming that explains this phenomenon. Consistent with that, Dirkx and colleagues reported previously that transition through quiescence increases the parasite’s infectivity and transmission potential ^26^.

The 11 *L. donovani* lines studied here exhibited here a range of quiescence and persister phenotypes, which may be exploited further to probe the molecular mechanism that control the transition to, maintenance and exit from quiescent state. For example, BPK087 did not exhibit reduction of rEGFP expression in the very early timepoints of exposure, resulting in rapid killing. This suggest that this line was not able to enter quiescence in our experimental conditions. It would be interesting to see whether this remains the same if different doses of PAT are applied. If BPK087 proves to be unable to enter into quiescence in response to PAT pressure, it could serve as a useful model to uncover signalling that leads to quiescence entry. At the same time, several lines characterized by profile 3 exhibited the ability to exit quiescence still under drug pressure and could be employed to uncover the mechanisms controlling the maintenance and promoting the exit of quiescent state.

This study was carried out with the extracellular promastigote form of *Leishmania* and should be repeated with both axenic and intracellular amastigote forms to confirm the same phenomena are applicable to the clinically relevant stage. Several studies have already indicated the existence of persister-like cell in *Leishmania* infections of mammals and it seems increasingly likely that *L. donovani* persister-like cells can explain some cases of treatment failure in the absence of drug resistance and post-kala azar dermal leishmaniasis (PKDL), observed years after clinical healing of VL.

In summary, this report demonstrates a worrying, intrinsic ability of *Leishmania* to tolerate high lethal doses of a drug and exhibit cross-tolerance to different drugs, even with non-overlapping modes of action. We also unravel unexpected effects of adoption of quiescence, which may go beyond the period of exposure to drug. In strains with pre-existing genomic adaptations to the drug, transition thorough the quiescent state potentially facilitates the development of a form of population-wide super drug resistance. Thus, even a short period of quiescence may have much more serious impact than previously thought, with a potential to create super-resistant parasites.

## Methods

### Parasites

Eleven strains of *L. donovani* were used in this study (Table 01). Ten were derived from clinical isolates originating from VL patients in Nepal and classed into 3 main genotypes according to the amplification or not of H-locus (containing MRPA) and AQP1 sequence ^16^ (i) No amplification of H-locus and wild type AQP1 (H-/AQP+): MHOM/NP/2002/BPK026/0 clone 5 (LdBPK026), MHOM/NP/2002/BPK031/0 clone 2 (LdBPK031) and MHOM/NP/2002/BPK156/0 (LdBPK156). **(ii)** Amplification of H-locus and wild type AQP1 (H+/AQP+): MHOM/NP/2002/BPK080/0 clone 1 (LdBPK080), MHOM/NP/2002/BPK085/0 clone 8 (LdBPK085), MHOM/NP/2002/BPK087/0 clone 11 (LdBPK087), MHOM/NP/--/BPK190/0 clone 3 (LdBPK190), MHOM/NP/2003/BPK282/0 clone 4 (LdBPK282), MHOM/NP/2003/BPK294 clone 1 (LdBPK294). **(iii)** Amplification of H-locus and AQP1 indel in position Ld31_0007735 (H+/AQP-): MHOM/NP/2003/BPK275/0 clone 18 (LdBPK275). A last strain originating from Sudan was used as a reference: MHOM/Sudan/1962/1S-2D Bob (LdBOB). All strains were cryopreserved at the Institute of Tropical Medicine in Antwerp, Belgium. Promastigotes were maintained at 26°C in M199 medium (Thermo Fisher) at pH 7.2 supplemented with 20% of heat-inactivated fetal bovine serum (HI-FBS), 5 mg/mL hemin, 10µM acid folic, 1X RPMI vitamin mix, 2mM glutamine, 100 units/mL of penicillin, and 100 µg/mL of streptomycin, which was referred to as M199-sup in the following experiments. For transfections, HOMEM medium (Gibco) supplemented with 20% HI-FBS, referred to as HOMEM-sup, was used to transfect *Leishmania* promastigotes mentioned below.

### Drugs

The drugs used in this study were: i) for leishmaniosis treatment: potassium antimonyl tartrate, (383376, Sigma-Aldrich), miltefosine (M5571, Sigma-Aldrich), amphotericin B (A2942, Sigma-Aldrich), and paromomycin (P5057, Sigma-Aldrich); ii) in development: bortezomib (5.04314, Sigma-Aldrich), and carbonyl cyanide 4-(trifluoromethoxy) phenylhydrazone (C2920, Sigma-Aldrich). The stock solutions of potassium antimonyl tartrate (PAT), paromomycin (PARO), and miltefosine (MIL) were dissolved in PBS, meanwhile bortezomib (BOR), and carbonyl cyanide 4-(trifluoromethoxy) phenylhydrazone (FCCP) in dimethylsulfoxide (DMSO) at a concentration of 20-50 mM and stored at - 20°C. The working solutions were freshly diluted in PBS or dimethylsulfoxide (DMSO) and added to M199-sup until reaching the desired final concentrations for the different drugs/inhibitors. A control containing either PBS (0.5% v/v) or DMSO (0.1% v/v), depending on which was used to dissolve the drug or inhibitor, was included in the plate. The addition of PBS or DMSO did not interfere with the parasite growth.

### Engineering of fluorescent lines

For each strain, the enhanced green fluorescent protein (EGFP) was integrated within the 18S ribosomal DNA locus (rEGFP) of promastigotes, with pLEXSY-hyg system (Jena Bioscience). rEGFP was previously validated as a negative marker for quiescence: highly expressed in proliferative cells and downregulated during quiescence ^36^. For transfection, *L. donovani* promastigotes were maintained at 26°C in HOMEM-sup with passages each 4-7 days at a dilution 1/50. A quantity of 2 × 10⁷ promastigotes was transfected with 100 ng of the linearized DNA fragment containing EGFP, using the Basic Parasite Nucleofector kit 1 (VMI 1011, Lonza); program U-033. After 24 hours of transfection, promastigotes harboring rEGFP were selected with hygromycin B at a final 25 µg/ml concentration. Parasites were passaged 3 times in HOMEM-sup with hygromycin B until the isolation of individual clones. An appropriate dilution of 2-day promastigotes culture was washed with PBS and adjusted to a concentration of 2 × 10^6^ parasites/mL in PBS containing 2% HI-FBS, 100 units/mL of penicillin, and 100 µg/mL of streptomycin. Then, the suspension was filtered using a 5-µM pore filter to remove cell clumps. A total of 8 individual clones were sorted by fluorescence-activated cell sorting (FACS), using S3e TM Cell Sorter (BIORAD) using a 488 nm laser for EGFP excitation. Single cells expressing EGFP were selected based on FSC-A vs SSC-A followed by FSC-A vs FSC-H, and FSC-A vs EGFP, and collected into 300 µL of HOMEM-sup. The clonal expansion was monitored for two weeks until enough promastigotes were present for further passages. The EGFP fluorescence of the clones was assessed by FACSVerse cytometer (BD Biosciences), where the clones with more than 99% of cells expressing EGFP and the highest median fluorescence intensity (MFI) were selected for further experiments. The generated lines were called rEGFP lines, maintained in M199-sup, and then cryo-preserved.

### Estimation of PAT concentration at which parasites reach a minimal metabolic activity

Promastigotes of the *L. donovani* rEGFP lines were synchronized by consecutive passages every two days to ensure that the promastigotes were in the proliferative phase. Then, parasites were distributed (5×10⁵ parasites/mL) in 200µL of M199-sup in sterile 96-well culture plates and exposed to different concentrations of PAT ranging from 1200 to 12.5 µM for 4 days at 26°C. An additional well was included with only M199-sup as a control for each *L. donovani* line. Resazurin sodium salt (Sigma) was added at a final concentration of 80µM, and incubated for 5 h in the dark at 26°C. The fluorescence was measured using the Victor X2 Multilaber Reader (PerkinElmer) with a 560 excitation and 590 emission wavelength. The fluorescence measurements for each line were normalized in comparison to their respective control with only M199-sup to calculate the percentage of cells with metabolic activity. The lethal PAT concentrations for each line were determined as the drug concentration point where the percentage of metabolic active cells was less ∼1%.

### Monitoring quiescence and cell viability during PAT exposure

The rEGFP lines were passaged twice each two days to synchronize them at the logarithmic phase and then a density of 5×10⁵ parasites/ml was exposed to a normally lethal PAT doses during 10 days at 26°C where M199-sup with PAT was refreshed once after 5 days. The respective controls were prepared for each line with only M199-sup, and promastigotes were collected daily. At each time point, parasites were centrifugated at 1800 x g for 10 minutes, resuspended in PBS with 1 µM NucRed Dead 647 (Thermo Fisher), a dead marker that binds to DNA when the cell membrane is damaged and emits fluorescence. After 10 minutes at room temperature, the samples were analyzed by FACSVerse (BD Biosciences) cytometer including as controls: (i) negative control, wild-type line, and not-stained, and (ii) a dead control, parasites exposed to thermal shock at 70°C for 10 minutes. The gating scheme was initiated with a large gate of FSC-A vs SSC-A to include all cells due to the heterogeneous size and complexity of the parasites, continued with FSC-H vs FSC-A gate to avoid doublets or aggregates. A gate of FSC-A vs NucRed Dead (−) was used to exclude dead cells, and live cells were determined based on a gate of NucRed Dead (−) vs EGFP(+). The Flow Cytometry Standard (FCS) data files were analyzed with the FlowJo™ v10.10 to determine the median fluorescence intensity (MFI) of EGFP and the percentage of viable cells.

### Reversibility test

Promastigotes from the different *L. donovani* rEGFP lines under PAT exposure during all the combination and serial conditions were harvested and washed once with M199 medium by centrifugation at 1800 x g for 10 minutes. The supernatant was removed without disturbing the pellet, resuspended in M199-sup, and incubated at 26°C. Then, we monitored the cultures weekly for 21 days under microscopic observation until flagellated mobile promastigotes were observed.

### Promastigote susceptibility assay

Susceptibility of the promastigotes was assessed in the eight lines that survive PAT exposure and revert to a proliferative state, including their respective controls maintained in only M199-sup medium. Likewise, the promastigotes of BPK026 and BPK275 were used to measure the susceptibility against five drugs for the cross-tolerance experiments. The measurement was performed using the resazurin method as previously described with minor modifications ^20,50^. Briefly, logarithmic-stage promastigotes (5×10⁵ parasites/mL) were plated in 200µL of M199-sup in sterile 96-well culture plates and grown in the presence of M199-sup alone (control) and medium containing serial dilution of drugs for a period of 72 h at 26°C. Then resazurin sodium salt (Sigma) was added at a final concentration of 80 µM, and the parasites were incubated for 5 h at 26°C protected from light. The fluorescence was recorded with an excitation/emission 560/590 nm using the Victor X2 Multilabel Reader (PerkinElmer). All experiments were done with three independent cultures with two or three technical replicates each. The IC_50_ (50% inhibitory concentration measurement) was calculated with GraphPad Prism 8 using a sigmoidal dose-response model with variable slope.

### DNA preparation and bulk whole-genome sequencing (WGS) analysis

Promastigotes were harvested from the eight lines that reverted to a proliferative state after PAT exposure or controls maintained without it. The parasites were washed with PBS and the genomic DNA was extracted using the QIAamp DNA Mini Kit (51306, Qiagen) following the manufacturer’s recommendations. The DNA concentration was measured using the Qubit dsDNA Quantification Assay Kit broad-range (Q32853, Thermo Fisher). The libraries PCR-free were prepared and sequenced on the Illumina NovaSeq6000 platform using 2 x 150 bp paired-end reads aiming for a sequence depth of ∼100x for each sample, a service provided by GenomeScan (Netherlands).

After quality control of the raw fastq files using FastQC, the reads were mapped to the *L. donovani* reference genome LdBPK282-V2 ^19^, using BWA v0.7.17 using a seed length of 50. Reads with a mapping quality higher than 30 were selected using SAMtools, duplicate reads removed using the Picard MarkDuplicates command (version 2.22.4), and only properly paired reads were retained using samtools. The somy values were estimated as the median reads count per million (CPM) per 10kb bin of chromosomes normalized by the median CPM per bin of 5 chromosomes surrounding on both sides. SNP calling was conducted using the Genome Analysis Toolkit (GATK) pipeline (version 4.1.4.1) following GATK’s best practices guidelines. The process included: (1) generating GVCF-formatted files using GATK HaplotypeCaller, (2) combining the GVCF files from all samples with the GATK CombineGVCF command, (3) performing genotyping with the GATK GenotypeGVCF command, and (4) filtering SNPs and indels based on GATK’s recommended best practices using the SelectVariants and VariantFiltration commands. For the CNV detection, for every sample the sequencing depth per position was derived using the SAMtools depth command. The CNV value was calculated by taking the ration of the median coverage per gene over the median coverage of the chromosome the gene is located on. Statistical tests comparing either CNV or Somy values were performed using a student’s t-test. Visualization of somy values, SNP data and CNV values was performed in R (version 4.2.0) using heatmaps and dot plots. Heatmaps were created with the pheatmap package (version 1.0.12), while dot plots were generated using the ggplot2 package (version 3.4.1).

### Determination of mitochondrial membrane potential (MMP)

The BPK026 and BPK275 promastigotes were synchronized by two-day passaging in M199-sup before each experiment. Several cultures were prepared with a density of 5×10^5^ parasites/mL with normally lethal doses of PAT (50 µM for BPK026 and 200 µM for BPK275) at different times to have samples under drug exposure for 2, 4, and 10 days with their respective controls with only M199-sup. The mitochondrial membrane potential was determined using the MitoProbe™ DilC₁ (5) Assay Kit (M341151, Thermo Fisher) based on the manufacturer’s instructions. Parasites under PAT exposure were centrifuged at 1800 x g for 10 minutes, the pellet was resuspended in M199 medium and disaggregated into a single-cell suspension passing 6 times through a 26-G needle and syringe. Promastigotes were centrifuged at 1800 x g for 10 minutes, washed once in PBS, and resuspended at a concentration of 2 x10⁶ parasites/ml in PBS containing 100nM MitProbe DilC₁(5) for 45 minutes at 26°C. Then, 1 µM SYTOX Blue Dead Cell stain (S34857, Thermo Fisher) was added to label dead cells for 5 minutes. In the experiment three controls were used: (i) a dead control, heat-killed parasites at 70°C for 10 min, (ii) a positive control, parasites treated for 10 minutes with 200µM carbonyl cyanide m-clorophenylhydrazone (CCCP) which disrupts mitochondrial membrane potential, and (iii) a negative control, consisting of unstained parasites of a line without EGFP gene inserted. Also, fluorescence minus one (FMO) controls were used to assess autofluorescence and establish the gates for the selection of the cell populations. Subsequently, the samples were analyzed by FACSVerse cytometer (BD Biosciences) using lasers 405nm for SYTOX Blue dye, 480nm for EGFP, and 640nm for MitoProbe dye. Flow cytometric analysis was performed using FlowJo™ version 10.10.

### Proliferation analysis

Approximately 1.5 × 10^7^ promastigotes from *L. donovani* BPK026 and BPK275 were labeled with CellTracker Deep Red (C34565, Thermo Fisher technologies) following the manufacturer’s instructions. Briefly, parasites were collected, washed once with PBS, and resuspended in 1 mL of M199 medium containing 5 µM of CellTracker Deep Red. After 40 minutes of incubation at 26°C, cells were washed with M199 medium, centrifuged at 1500 x g for 10 minutes, and resuspended in pre-warmed M199-sup. Following labeling, 5 × 10^5^ promastigotes/mL were treated with PAT at 50 µM (BPK026) and 200 µM (BPK275) in M199-sup or only M199-sup as a control and incubated at 26°C protected from the light. The parasites were collected at three time points: 0, 4, and 10 days. At each measurement point, promastigotes under PAT exposure were disaggregated using a 26-G needle and syringe and centrifuged at 1800 x g for 10 minutes. The pellet and promastigotes from the controls were resuspended in PBS at a concentration of 2 × 10⁶ parasites/mL, and labeled with 1 µM SYTOX Blue Dead Cell stain (S34857, Thermo Fisher) for 5 minutes in the dark, as a cell death marker. The fluorescence data was acquired by FACSVerse cytometer (BD Biosciences) using lasers 405nm for SYTOX Blue dye, 480nm for EGFP, and 640nm for CellTracker Deep Red dye. Flow cytometric analysis was performed using FlowJo™ version 10.10.

### Cross-tolerance experiment

To evaluate tolerance to other anti-leishmanial drugs and drugs in development, of PAT-tolerant promastigotes, we exposed the promastigotes of BPK026 and BPK275 to 5 drugs with different mechanisms of action. We performed susceptibility tests by resazurin assay, mentioned above, to calculate the IC50 estimates. As the promastigotes were already under high stress using normally lethal PAT doses (50µM for BPK026 and 200µM for BPK275), we decided to use the IC50 estimates of each drug. We designed an experimental design consisting of two exposure times of 5 days each, for a total duration of 10 days where we had two conditions for each drug: (i) Combination, where the lethal concentration of PAT and the drug (IC50 estimate) were present in both exposure times, (ii) Serial, promastigotes were under normally lethal PAT concentration for the first exposure, followed by a mix of PAT with the other drug for the second exposure. As a control group, promastigotes were under only lethal PAT concentrations for both exposure times. The promastigotes were collected, centrifuged at 1800 x g for 10 minutes, and resuspended in M199 medium. The samples with large aggregates were passed through 26-G to have single-cell suspension. The parasites were centrifuged again and resuspended in PBS with NucRed Dead as a viability marker. The acquisition of the fluorescence data and its gating scheme for analysis was the same as mentioned above.

### Statistical analyses

Statistical analysis and data visualization were performed in R studio (R version 4.3.1) using Student’s t-test, with p-value < 0.05 considered statistically significant, and ANOVA followed by a post hoc Tukey’s multiple comparison test.

## Acknowledgments

This study received financial support from the Flemish Fund for Scientific Research grant G075121N. Isolation and characterization of clinical isolates was financed by the European Commission through Leishnatdrug-R project, grant ICA4-CT-2001-10076. M.A.D. acknowledges support from Flemish Ministry of Science and Innovation.

## Competing interests

The authors declare no competing interests.

## Additional information

Supplementary material is available in Aroni-Soto_supplement.pdf

## Notes

### Competing Interest Statement

The authors have declared no competing interest.

### Summary of Updates

In previously preprinted version, table 3 was missing. This has been corrected now

## References

1. WHO. Antimicrobial resistance. https://www.who.int/news-room/fact-sheets/detail/antimicrobial-resistance (2023).

2. Gow, N. A. R. et al. The importance of antimicrobial resistance in medical mycology. Nature Communications 2022 13:1 13, 1–12 (2022).

3. Darby, E. M. et al. Molecular mechanisms of antibiotic resistance revisited. Nature Reviews Microbiology 2022 21:5 21, 280–295 (2022).

4. Balaban, N. Q. et al. Definitions and guidelines for research on antibiotic persistence. Nature Reviews Microbiology 2019 17:7 17, 441–448 (2019).

5. Dewachter, L., Fauvart, M. & Michiels, J. Bacterial Heterogeneity and Antibiotic Survival: Understanding and Combatting Persistence and Heteroresistance. Mol Cell 76, 255–267 (2019).

6. Peyrusson, F. et al. Intracellular Staphylococcus aureus persisters upon antibiotic exposure. Nature Communications 2020 11:1 11, 1–14 (2020).

7. Nguyen, D. et al. Active starvation responses mediate antibiotic tolerance in biofilms and nutrient-limited bacteria. Science (1979) 334, 982–986 (2011).

8. Keren, I., Shah, D., Spoering, A., Kaldalu, N. & Lewis, K. Specialized persister cells and the mechanism of multidrug tolerance in Escherichia coli. J Bacteriol 186, 8172–8180 (2004).

9. Rittershaus, E. S. C., Baek, S. H. & Sassetti, C. M. The Normalcy of Dormancy: Common Themes in Microbial Quiescence. Cell Host Microbe 13, 643–651 (2013).

10. Sun, S. & Gresham, D. Cellular quiescence in budding yeast. Yeast 38, 12–29 (2021).

11. Amato, S. M. et al. The role of metabolism in bacterial persistence. Front Microbiol 5, 79557 (2014).

12. Ponte-Sucre, A. et al. Drug resistance and treatment failure in leishmaniasis: A 21st century challenge. PLoS Negl Trop Dis 11, e0006052 (2017).

13. Hefnawy, A., Berg, M., Dujardin, J. C. & De Muylder, G. Exploiting Knowledge on Leishmania Drug Resistance to Support the Quest for New Drugs. Trends Parasitol 33, 162–174 (2017).

14. Mandal, S., Maharjan, M., Singh, S., Chatterjee, M. & Madhubala, R. Assessing aquaglyceroporin gene status and expression profile in antimony-susceptible and -resistant clinical isolates of Leishmania donovani from India. Journal of Antimicrobial Chemotherapy 65, 496–507 (2010).

15. Potvin, J. E., Leprohon, P., Queffeulou, M., Sundar, S. & Ouellette, M. Mutations in an Aquaglyceroporin as a Proven Marker of Antimony Clinical Resistance in the Parasite Leishmania donovani. Clinical Infectious Diseases 72, e526–e532 (2021).

16. Imamura, H. et al. Evolutionary genomics of epidemic visceral leishmaniasis in the Indian subcontinent. Elife 5, (2016).

17. Dumetz, F., et al. Molecular Preadaptation to Antimony Resistance in Leishmania donovani on the Indian Subcontinent. mSphere 3, (2018).

18. Downing, T. et al. Whole genome sequencing of multiple Leishmania donovani clinical isolates provides insights into population structure and mechanisms of drug resistance. Genome Res 21, 2143–2156 (2011).

19. Dumetz, F. et al. Modulation of aneuploidy in leishmania donovani during adaptation to different in vitro and in vivo environments and its impact on gene expression. mBio 8, (2017).

20. Hefnawy, A. et al. Genomic and Phenotypic Characterization of Experimentally Selected Resistant Leishmania donovani Reveals a Role for Dynamin-1-Like Protein in the Mechanism of Resistance to a Novel Antileishmanial Compound. mBio 13, (2022).

21. Shaw, C. D. et al. In vitro selection of miltefosine resistance in promastigotes of Leishmania donovani from Nepal: genomic and metabolomic characterization. Mol Microbiol 99, 1134–1148 (2016).

22. Barrett, M. P., Kyle, D. E., Sibley, L. D., Radke, J. B. & Tarleton, R. L. Protozoan persister-like cells and drug treatment failure. Nature Reviews Microbiology 2019 17:10 17, 607–620 (2019).

23. Jara, M. et al. Unveiling drug-tolerant and persister-like cells in Leishmania braziliensis lines derived from patients with cutaneous leishmaniasis. Front Cell Infect Microbiol 13, 1253033 (2023).

24. Jara, M. et al. Transcriptional Shift and Metabolic Adaptations during Leishmania Quiescence Using Stationary Phase and Drug Pressure as Models. Microorganisms 10, 97 (2022).

25. Kloehn, J. et al. Identification of metabolically quiescent leishmania mexicana parasites in peripheral and cured dermal granulomas using stable isotope tracing imaging mass spectrometry. mBio 12, (2021).

26. Dirkx, L. et al. Long-term hematopoietic stem cells trigger quiescence in Leishmania parasites. PLoS Pathog 20, e1012181 (2024).

27. Kloehn, J., Saunders, E. C., O’Callaghan, S., Dagley, M. J. & McConville, M. J. Characterization of Metabolically Quiescent Leishmania Parasites in Murine Lesions Using Heavy Water Labeling. PLoS Pathog 11, e1004683 (2015).

28. Mandell, M. A. & Beverley, S. M. Continual renewal and replication of persistent Leishmania major parasites in concomitantly immune hosts. Proc Natl Acad Sci U S A 114, E801–E810 (2017).

29. Dirkx, L. et al. Long-term hematopoietic stem cells as a parasite niche during treatment failure in visceral leishmaniasis. Communications Biology 2022 5:1 5, 1–15 (2022).

30. Brotherton, M. C. et al. Proteomic and Genomic Analyses of Antimony Resistant Leishmania infantum Mutant. PLoS One 8, e81899 (2013).

31. Patino, L. H. et al. Major changes in chromosomal somy, gene expression and gene dosage driven by SbIII in Leishmania braziliensis and Leishmania panamensis. Scientific Reports 2019 9:1 9, 1–13 (2019).

32. Negreira, G. H. et al. High throughput single-cell genome sequencing gives insights into the generation and evolution of mosaic aneuploidy in Leishmania donovani. Nucleic Acids Res 50, 293–305 (2022).

33. Sterkers, Y. et al. Novel insights into genome plasticity in Eukaryotes: mosaic aneuploidy in Leishmania. Mol Microbiol 86, 15–23 (2012).

34. Wyllie, S., Cunningham, M. L. & Fairlamb, A. H. Dual Action of Antimonial Drugs on Thiol Redox Metabolism in the Human Pathogen Leishmania donovani. Journal of Biological Chemistry 279, 39925–39932 (2004).

35. Goyard, S. et al. An in vitro system for developmental and genetic studies of Leishmania donovani phosphoglycans. Mol Biochem Parasitol 130, 31–42 (2003).

36. Jara, M. et al. Tracking of quiescence in Leishmania by quantifying the expression of GFP in the ribosomal DNA locus. Scientific Reports 2019 9:1 9, 1–12 (2019).

37. Taneja, N. K. & Tyagi, J. S. Resazurin reduction assays for screening of anti-tubercular compounds against dormant and actively growing Mycobacterium tuberculosis, Mycobacterium bovis BCG and Mycobacterium smegmatis. Journal of Antimicrobial Chemotherapy 60, 288–293 (2007).

38. Chawla, B., Jhingran, A., Panigrahi, A., Stuart, K. D. & Madhubala, R. Paromomycin Affects Translation and Vesicle-Mediated Trafficking as Revealed by Proteomics of Paromomycin – Susceptible –Resistant Leishmania donovani. PLoS One 6, e26660 (2011).

39. Rakotomanga, M., Blanc, S., Gaudin, K., Chaminade, P. & Loiseau, P. M. Miltefosine Affects Lipid Metabolism in Leishmania donovani Promastigotes. Antimicrob Agents Chemother 51, 1425 (2007).

40. Stone, N. R. H., Bicanic, T., Salim, R. & Hope, W. Liposomal Amphotericin B (AmBisome®): A Review of the Pharmacokinetics, Pharmacodynamics, Clinical Experience and Future Directions. Drugs 2016 76:4 76, 485–500 (2016).

41. López-Arencibia, A., Bethencourt-Estrella, C. J., San Nicolás-Hernández, D., Lorenzo-Morales, J. & Piñero, J. E. Anti-COVID Drugs (MMV COVID Box) as Leishmanicidal Agents: Unveiling New Therapeutic Horizons. Pharmaceuticals 17, 266 (2024).

42. Sogbein, O. et al. Bortezomib in cancer therapy: Mechanisms, side effects, and future proteasome inhibitors. Life Sci 358, 123125 (2024).

43. Brauner, A., Fridman, O., Gefen, O. & Balaban, N. Q. Distinguishing between resistance, tolerance and persistence to antibiotic treatment. Nature Reviews Microbiology 2016 14:5 14, 320–330 (2016).

44. Balaban, N. Q., Merrin, J., Chait, R., Kowalik, L. & Leibler, S. Bacterial persistence as a phenotypic switch. Science (1979) 305, 1622–1625 (2004).

45. El Fadili, K., et al. Role of the ABC transporter MRPA (PGPA) in antimony resistance in Leishmania infantum axenic and intracellular amastigotes. Antimicrob Agents Chemother 49, 1988–1993 (2005).

46. Sun, L. L. et al. Aneuploidy enables cross-tolerance to unrelated antifungal drugs in Candida parapsilosis. Front Microbiol 14, 1137083 (2023).

47. Yang, F. et al. Aneuploidy Enables Cross-Adaptation to Unrelated Drugs. Mol Biol Evol 36, 1768–1782 (2019).

48. Negreira, G. H. et al. The adaptive roles of aneuploidy and polyclonality in Leishmania in response to environmental stress. EMBO Rep 24, (2023).

49. Sterkers, Y., Lachaud, L., Crobu, L., Bastien, P. & Pagès, M. FISH analysis reveals aneuploidy and continual generation of chromosomal mosaicism in Leishmania major. Cell Microbiol 13, 274–283 (2011).

50. Kulshrestha, A. et al. Validation of a simple resazurin-based promastigote assay for the routine monitoring of miltefosine susceptibility in clinical isolates of Leishmania donovani. Parasitol Res 112, 825–828 (2013).

