## supplementary information for "Multi-drug tolerance in *Leishmania* persister-like cells"

### SUPPLEMENTARY TABLES

**Supplementary Table 1.** Lowest PAT concentration at which no metabolic activity could be detected among promastigotes (see also Supp. Fig. 1), also called normally lethal dose.

| Strain | PAT µM |
|---|---|
| **P1** | |
| BPK026 | 50 |
| BPK031 | 100 |
| BPK156 | 200 |
| **P2** | |
| BPK190 | 400 |
| BPK087 | 400 |
| LdBOB | 400 |
| **P3** | |
| BPK085 | 400 |
| BPK282 | 600 |
| BPK294 | 400 |
| **P4** | |
| BPK080 | 400 |
| BPK275 | 200 |

**Supplementary Table 2.** Evolution of the viability percentage of promastigotes during 10 days of exposure to strain-specific normally lethal PAT doses. The values are represented as the mean of 3 biological replicates ± SEM.

| Strain | Viability (%) | | | | | | | |
|---|---|---|---|---|---|---|---|---|
| | D0 | D1 | D2 | D3 | D4 | D6 | D8 | D10 |
| **P1** | | | | | | | | |
| BPK026 | 100 ± 0 | 94.8 ± 0.1 | 73.2 ± 2.4 | 25.1 ± 1.6 | 5.1 ± 0.5 | 0.7 ± 0.2 | 0.5 ± 0.4 | 0.5 ± 0.2 |
| BPK031 | 100 ± 0 | 92 ± 1.6 | 44.6 ± 1.2 | 11.1 ± 1 | 5 ± 0.3 | 1.3 ± 0.5 | 0.6 ± 0.3 | 0.5 ± 0.1 |
| BPK156 | 100 ± 0 | 69.5 ± 5.9 | 51.4 ± 9.8 | 43.6 ± 7.4 | 22.7 ± 2.8 | 3.4 ± 1.3 | 0.6 ± 0.1 | 0.1 ± 0.1 |
| **P2** | | | | | | | | |
| BPK190 | 100 ± 0 | 58.2 ± 5.9 | 6 ± 1.6 | 1.4 ± 0.9 | 0.8 ± 0.5 | 0 ± 0 | 0 ± 0 | 0.5 ± 0.1 |
| BPK087 | 100 ± 0 | 57 ± 4.5 | 11 ± 0.6 | 2.4 ± 0.3 | 0.6 ± 0.2 | 0.1 ± 0.1 | 0 ± 0 | 0.5 ± 0.3 |
| LdBOB | 100 ± 0 | 22.9 ± 0.2 | 1.8 ± 0.3 | 5.5 ± 0.9 | 0.1 ± 0.1 | 0 ± 0 | 0 ± 0 | 0 ± 0 |
| **P3** | | | | | | | | |
| BPK085 | 100 ± 0 | 84.9 ± 2.6 | 35.9 ± 6.6 | 6.8 ± 1 | 1.9 ± 0.2 | 0.4 ± 0.3 | 0.8 ± 0.4 | 5.5 ± 3.3 |
| BPK282 | 100 ± 0 | 60 ± 0.4 | 15 ± 1.5 | 4.5 ± 0.9 | 3.2 ± 0.3 | 0.7 ± 0.2 | 0.9 ± 0.4 | 14.9 ± 4.5 |
| BPK294 | 100 ± 0 | 82.8 ± 2.4 | 39.1 ± 6.8 | 0.4 ± 0.2 | 3.4 ± 0.8 | 0.2 ± 0.1 | 0.1 ± 0.1 | 3.1 ± 0.6 |
| **P4** | | | | | | | | |
| BPK080 | 100 ± 0 | 85.3 ± 1.9 | 68.5 ± 6.2 | 54.4 ± 11.3 | 36.8 ± 14.1 | 10.2 ± 3.2 | 20.2 ± 7.4 | 53.1 ± 8.6 |
| BPK275 | 100 ± 0 | 92.1 ± 0.8 | 83 ± 0.9 | 75.7 ± 0.9 | 69.7 ± 1.4 | 80.1 ± 2.3 | 82.2 ± 1.5 | 92.3 ± 0.5 |

**Supplementary Table 3.** Reactivation of parasite proliferation (numbers out of 3 replicates) after PAT exposure for 4 and 10 days, at strain-specific normally lethal dose.

| Strain | H-locus amplified | AQP1 indel† | Reactivate after 4 day | Reactivate after 10 day |
|---|---|---|---|---|
| **P1** | | | | |
| BPK026 | no | no | 3 of 3 | 3 of 3 |
| BPK031 | no | no | 3 of 3 | 3 of 3 |
| BPK156 | no | no | 3 of 3 | 3 of 3 |
| **P2** | | | | |
| BPK190 | yes | no | 3 of 3 | 0 of 3 |
| BPK087 | yes | no | 3 of 3 | 1 of 3 |
| LdBOB | no | no | 3 of 3 | 0 of 3 |
| **P3** | | | | |
| BPK085 | yes | no | 3 of 3 | 3 of 3 |
| BPK282 | yes | no | 3 of 3 | 3 of 3 |
| BPK294 | yes | no | 3 of 3 | 3 of 3 |
| **P4** | | | | |
| BPK080 | yes | no | 3 of 3 | 3 of 3 |
| BPK275 | yes | yes | 3 of 3 | 3 of 3 |

† Indel associated with antimonial resistance.

**Supplementary Table 4.** Copy values H-locus (per haploid genome) of each *L. donovani* line after removal of PAT exposure (Post PAT), and controls without exposure (No PAT). The table shows the mean of three biological replicates ± SEM.

| Strain | H-locus amplified | H-locus copy values No PAT | H-locus copy values Post PAT |
|---|---|---|---|
| **P1** | | | |
| BPK026 | no | 0.88 ± 0.01 | 0.93 ± 0.03 |
| BPK031 | no | 0.93 ± 0.01 | 0.90 ± 0.01 |
| BPK156 | no | 0.95 ± 0.01 | 0.94 ± 0.01 |
| **P3** | | | |
| BPK085 | yes | 2.73 ± 0.03 | 2.78 ± 0.03 |
| BPK282 | yes | 2.75 ± 0.003 | 2.82 ± 0.09 |
| BPK294 | yes | 1.81 ± 0.04 | 1.84 ± 0.02 |
| **P4** | | | |
| BPK080 | yes | 1.91 ± 0.01 | 1.83 ± 0.03 |
| BPK275 | yes | 2.82 ± 0.03 | 2.69 ± 0.02 |

**Supplementary Table 5.** Susceptibility ($IC_{50}$) of BPK026 and BPK275 promastigotes to drugs with different modes of action: (i) used to treat leishmaniasis (MIL, Miltefosine; AmB, Amphotericin B, PARO, Paromomycin and (ii) in development (BOR, Bortezomib and FCCP (Carbonycil). Each value represents the mean of three biological replicates ± SEM.

| Strain | MIL IC50 µM | AMB IC50 nM | PARO IC50 µM | FCCP IC50 µM | BOR IC50 nM |
|---|---|---|---|---|---|
| **P1** | | | | | |
| BPK026 | 17.8 ± 0.1 | 64.2 ± 0.5 | 8.7 ± 0.1 | 20.0 ± 0.3 | 54.5 ± 0.6 |
| **P4** | | | | | |
| BPK275 | 13.0 ± 0.1 | 58.7 ± 0.7 | 21.4 ± 1.4 | 14.6 ± 0.5 | 55.5 ± 5.1 |

**Supplementary Table 6.** Experimental design for the study of multidrug tolerance, first exposure is made for 5 days, followed by a second exposure of another 5 days.

| | Drug/Inhibitor | First exposure | Second exposure |
|---|---|---|---|
| **Control** | PAT | PAT | PAT |
| **Combination** | AMB | PAT + AMB | PAT + AMB |
| **Serial** | | PAT | PAT + AMB |
| **Combination** | MIL | PAT + MIL | PAT + MIL |
| **Serial** | | PAT | PAT + MIL |
| **Combination** | FCCP | PAT + FCCP | PAT + FCCP |
| **Serial** | | PAT | PAT + FCCP |
| **Combination** | BOR | PAT + BOR | PAT + BOR |
| **Serial** | | PAT | PAT + BOR |
| **Combination** | PARO | PAT + PARO | PAT + PARO |
| **Serial** | | PAT | PAT + PARO |

**Supplementary Table 7.** Experimental design for the extended study of BOR tolerance. First exposure is made for 5 days, followed by a second exposure of another 5 days, PAT is used at a normally lethal dose; BOR is used at 1x, 2x and 5 x IC50.

| | Drug/Inhibitor | First exposure | Second exposure |
|---|---|---|---|
| **Control** | PAT | PAT | PAT |
| **Combination** | BOR | PAT + BOR | PAT + BOR |
| **Serial** | | PAT | PAT + BOR |
| **Combination** | 2 x BOR | PAT + BOR | PAT + BOR |
| **Serial** | | PAT | PAT + BOR |
| **Combination** | 5 x BOR | PAT + BOR | PAT + BOR |
| **Serial** | | PAT | PAT + BOR |

### SUPPLEMENTARY FIGURES

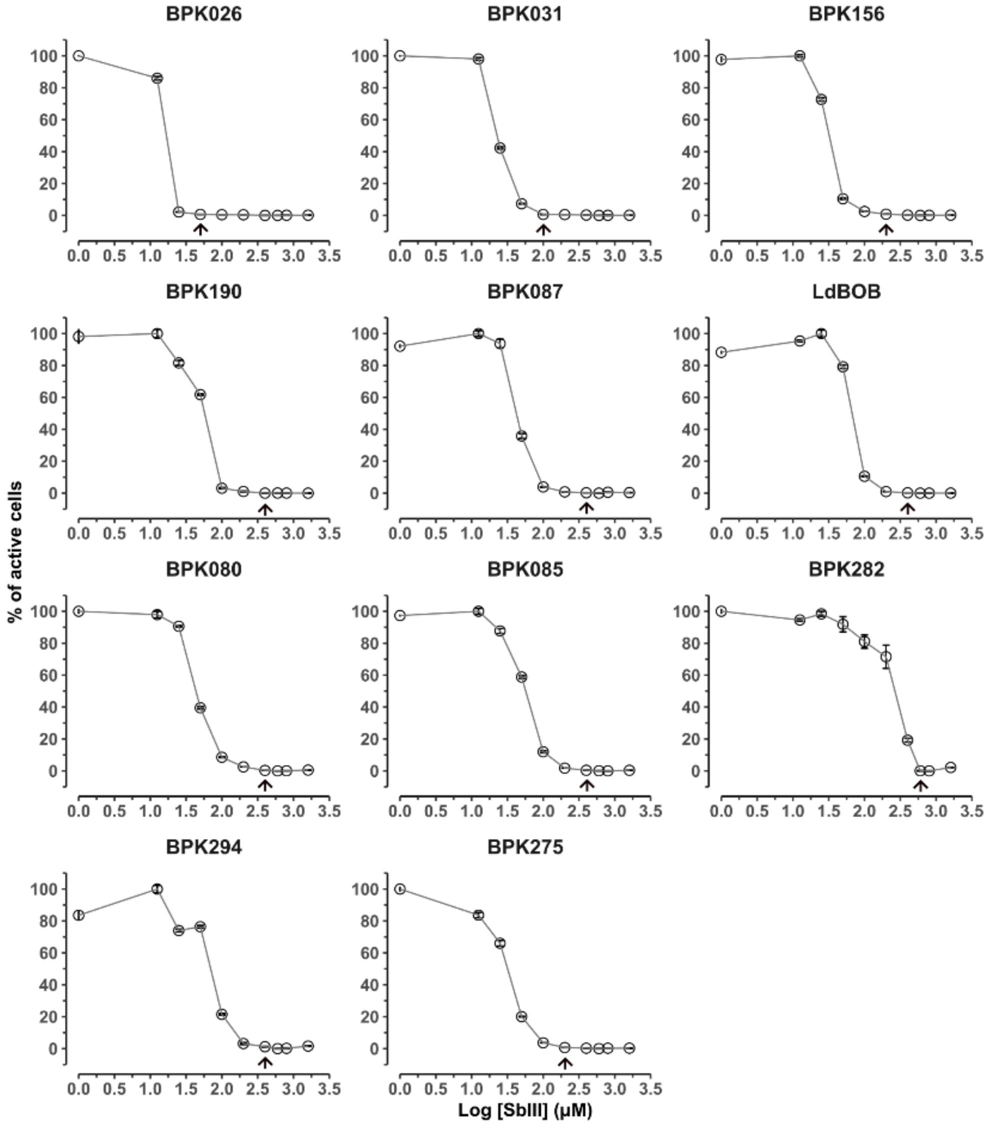


**Supplementary Figure 1.** Definition of the normally lethal dose of antimonials for each strain. Percentage of active cells in function of the PAT concentration measured to define the lowest PAT concentration (arrow) at which no metabolic activity could be detected (see also Supp. Table 1), further called normally lethal dose. Results are represented as the mean of 3 biological replicates ± SEM.

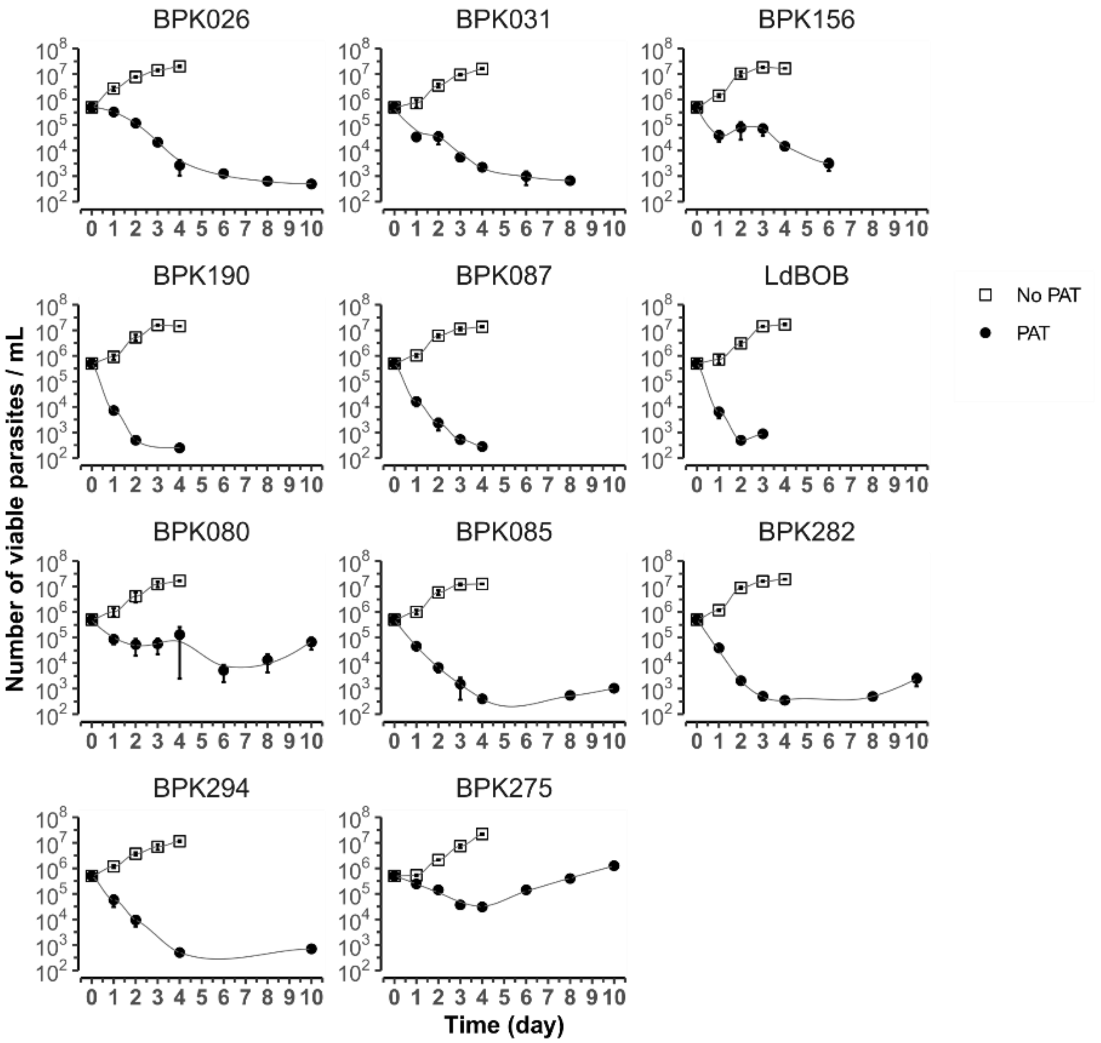


**Supplementary Figure 2. Growth curve of *L. donovani* strains in normally lethal PAT doses.** The number of viable promastigotes per mL during 10 days of PAT exposure (PAT), and a control of no drug exposure (No PAT) is included. The graph illustrates the mean ± SEM of three biological replicates for daily measurement.

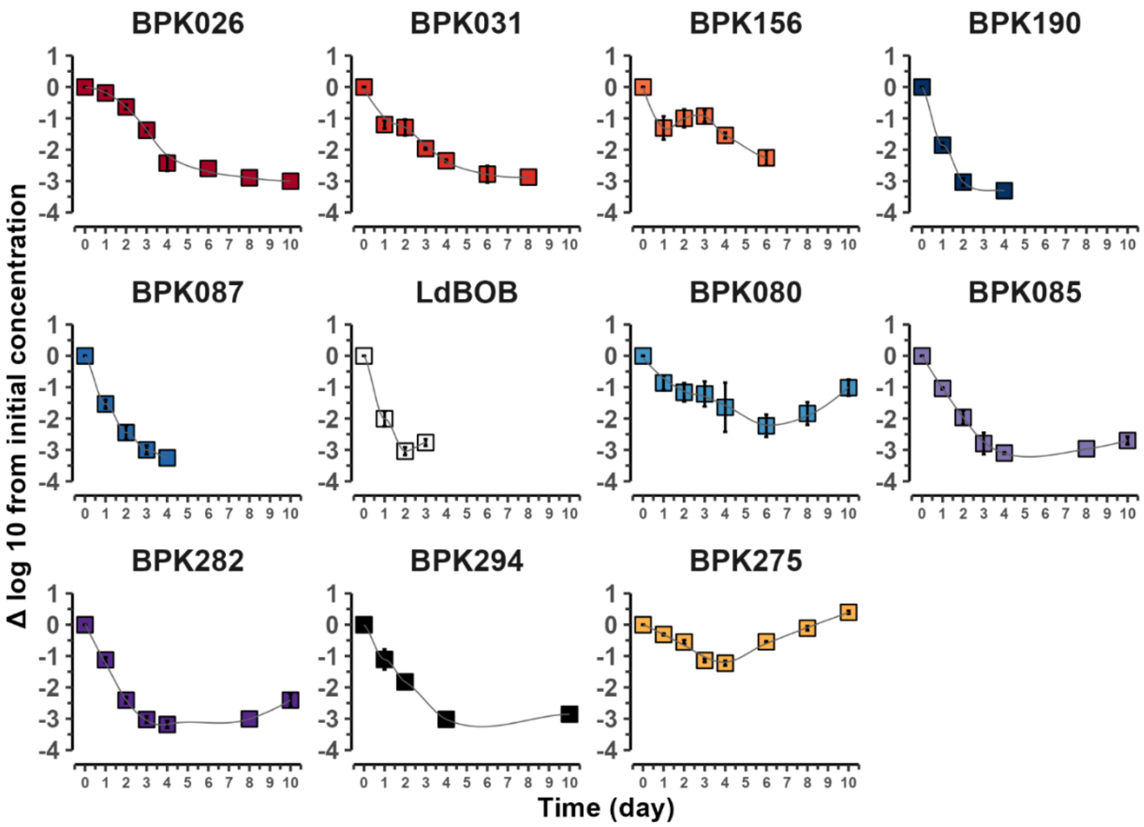


**Supplementary Figure 3. Survival of *L. donovani* strains under PAT pressure.** The number of viable promastigotes was measured by flow cytometry under 10-days exposure at strain-specific normally lethal doses of PAT. The values were calculated as the reduction in parasite concentration relative to the initial inoculum and are presented as the mean ± SEM of three biological replicates.

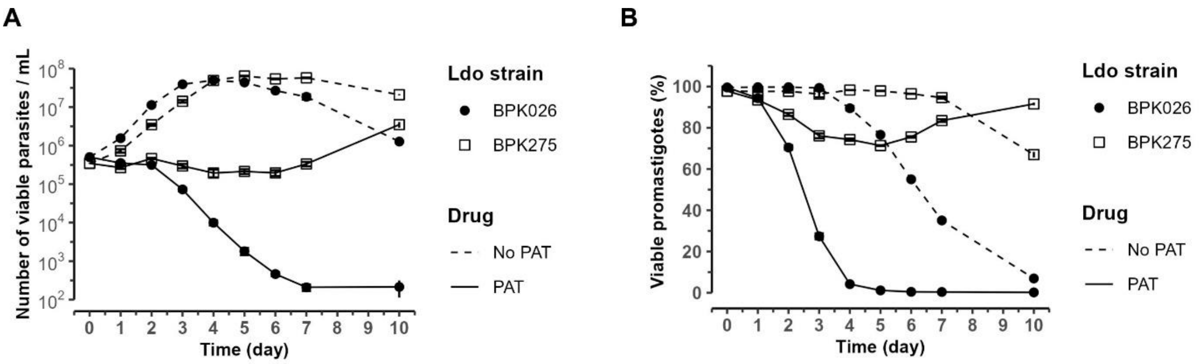


**Supplementary Figure 4. Growth and viability of promastigotes in normally lethal dose of PAT.** (**A**) Growth curve of promastigotes and, (**B**) Percentage of cell viability along 10 days of PAT exposure (PAT, solid line). A control of no drug exposure (No PAT, dashed line) is included. The data was acquired by flow cytometry for BPK026 (circle symbol) and BPK275 (square symbol). The graph illustrates the mean ± SEM of three biological replicates for daily measurement.

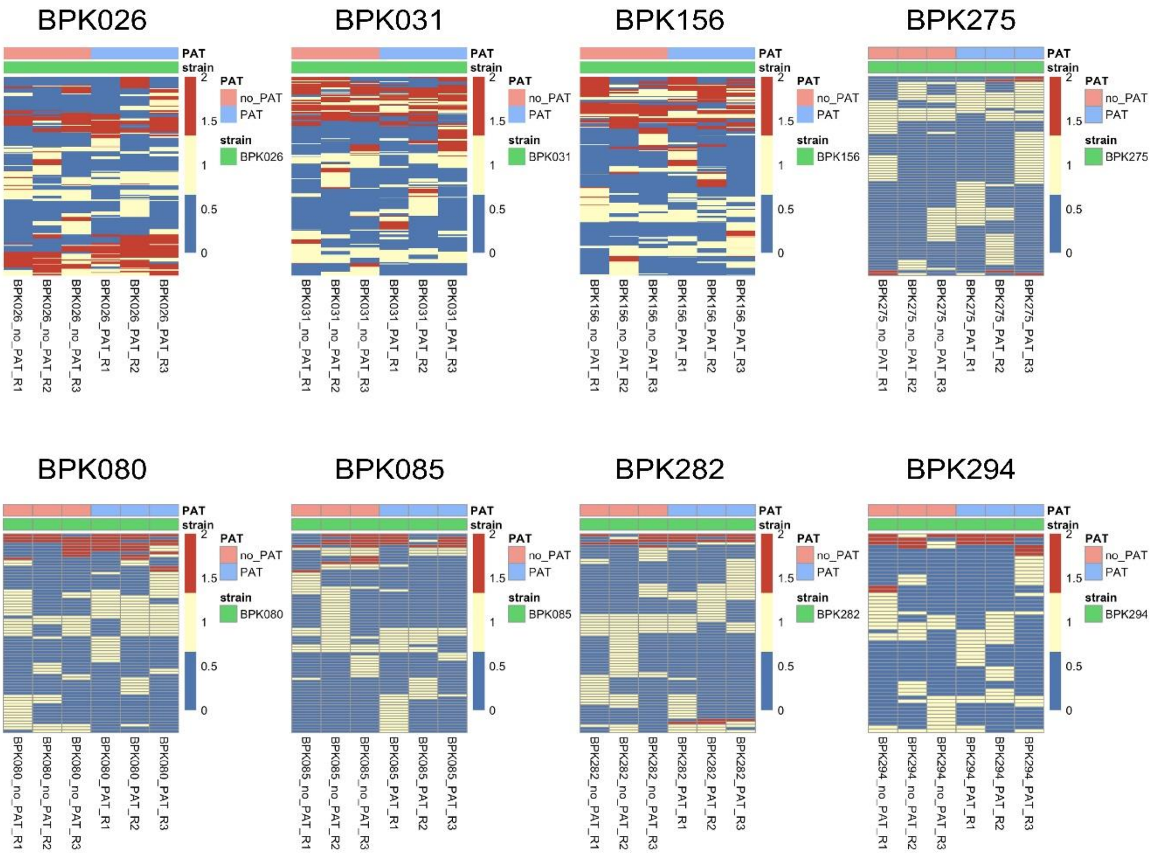


**Supplementary Figure 5. Heatmap illustrating the distribution of single nucleotide polymorphisms (SNPs) per individual strain.** SNPs that exhibited no variation across the six replicates per strain were filtered out. The color scheme categorizes SNPs as follows: blue represents the absence of a SNPs, yellow indicates heterozygous SNPs, and red denotes homozygous SNPs. Hierarchical clustering of samples (columns) and SNPs (rows) was performed using the complete linkage method with Euclidean distance. A color-coded bar above the heatmap indicates whether each strain was exposed to PAT, alongside the strain names. The graphs represents three biological replicates (R1, R2, R3) after removing PAT exposure, along with their corresponding controls without exposure (no_PAT).

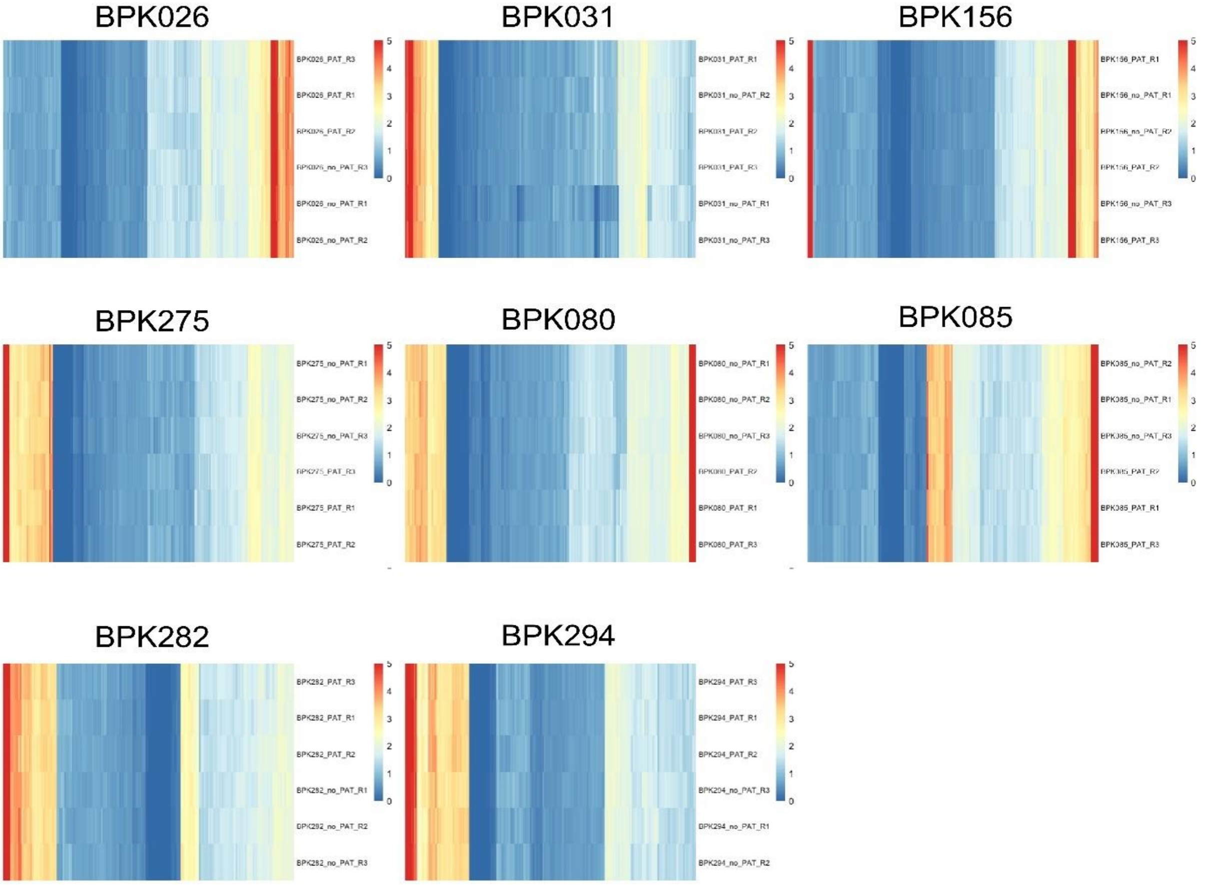


**Supplementary Figure 6. Heatmap for each strain** separately displaying CNV values for genes with a 50% increase or decrease in CNV (no statistical significance required).

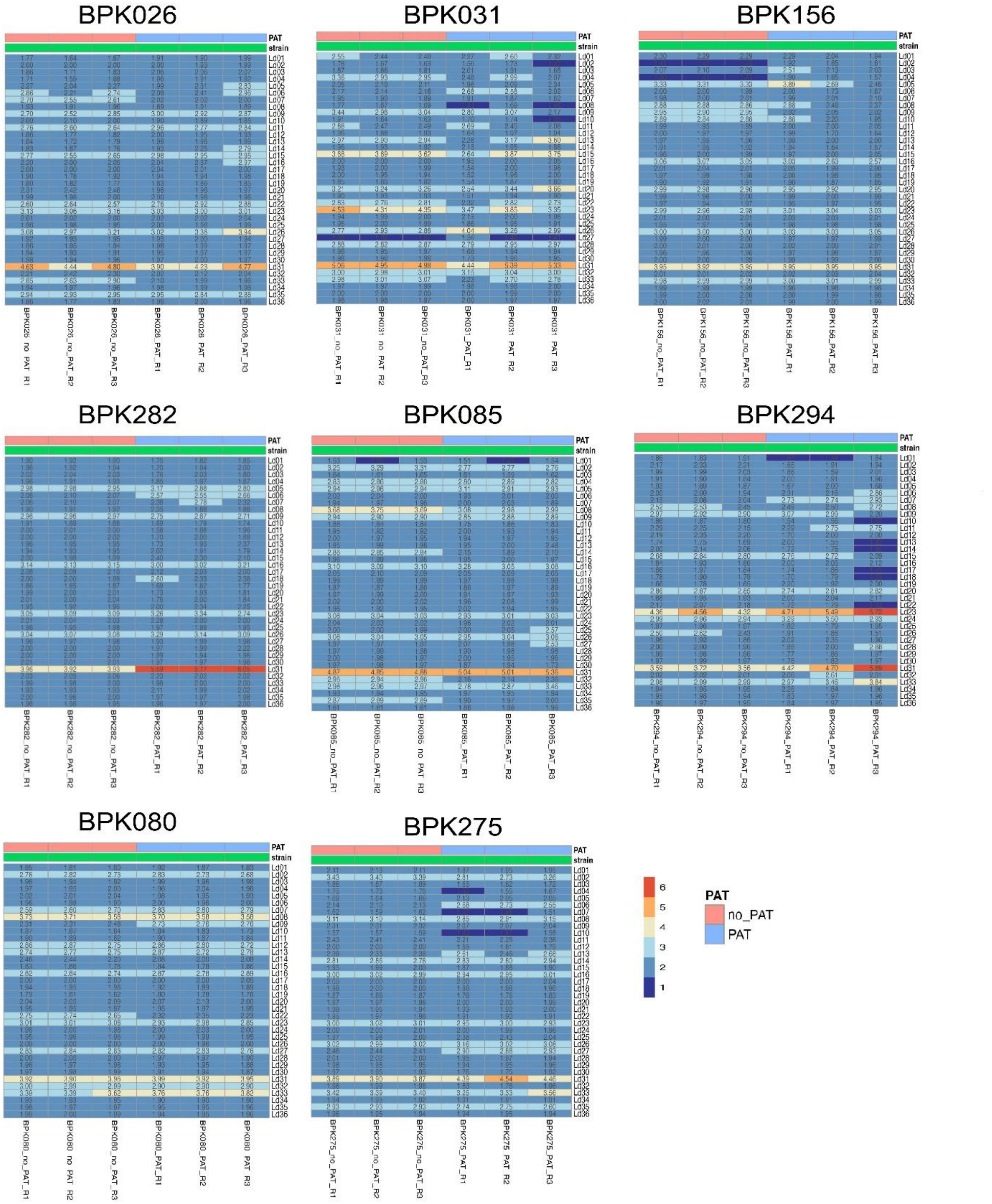


**Supplementary Figure 7. Heatmap showing the somy value per replicate, chromosome, and strain.** A color-coded bar above the heatmap indicates whether each strain was exposed to PAT. Three biological replicates are indicated as R1, R2, and R3.
